# A cortical output channel for perceptual categorization

**DOI:** 10.1101/2025.09.05.674544

**Authors:** Nathan A. Schneider, Michael I. Malina, Ross S. Williamson

**Affiliations:** Department of Otolaryngology, University of Pittsburgh, Pittsburgh, PA; Department of Neurobiology, University of Pittsburgh, Pittsburgh, PA; Department of Bioengineering, University of Pittsburgh, Pittsburgh, PA; Neuroscience Institute, Carnegie Mellon University, Pittsburgh, PA; Center for the Neural Basis of Cognition, Pittsburgh, PA; Pittsburgh Hearing Research Center, University of Pittsburgh, Pittsburgh, PA

## Abstract

Perceptual categorization allows the brain to transform diverse sensory inputs into discrete representations that support flexible behavior [1–7]. Auditory cortex (ACtx) has been implicated in this process [8–14], but the cell-type-specific circuits that implement category learning remain unknown. We trained head-fixed mice to categorize the temporal rate of amplitude-modulated noise while performing longitudinal two-photon imaging of layer (L)5 extratelencephalic (L5 ET) neurons alongside comparison populations of L2/3 and L5 intratelencephalic (L5 IT) neurons. With learning, L5 ET neurons underwent pronounced tuning modifications and developed robust, categorical responses, whereas L2/3 and L5 IT neurons did not. This categorical code was task engagement-dependent: it was present during behavior and absent during passive listening in the same neurons on the same day, indicating context-gated expression. Using a generalized linear model to dissociate stimulus- from choice-related signals, we confirmed that categorical selectivity in L5 ET neurons reflected sensory encoding rather than motor confounds. All three populations carried choice signals, but these were strongest in L5 ET neurons, suggesting a role in linking sensory categorization to action selection. These findings identify a projection-specific, deep-layer cortical output channel in which L5 ET neurons acquire categorical representations and selectively propagate behaviorally relevant signals to downstream targets.

## Introduction

Categorization is a fundamental aspect of perception, allowing organisms to compress a virtually infinite range of sensory inputs into a finite set of behaviorally meaningful groups [1–5, 15]. In audition, sounds with diverse acoustic features are mapped onto discrete perceptual labels that, in turn, guide decisions and actions [5–7, 16–21]. This capacity for auditory categorization is critical for survival, supporting predator avoidance, foraging, and communication [5, 22, 23], and underlies speech comprehension in humans [7]. To accomplish categorization, the auditory cortex (ACtx) must integrate and transform diverse spectrotemporal inputs into task-relevant representations. Neurophysiological studies demonstrate that ACtx neurons can represent auditory categories [8–14, 24], but the cell types and circuits involved in forming these representations and leveraging them to guide decisions remain poorly understood.

Complex behaviors emerge from coordinated interactions across distributed brain networks. Understanding auditory categorization therefore requires delineating how ACtx communicates with, and influences, other regions. ACtx is not homogeneous; it is organized into layers containing distinct excitatory cell types with characteristic projection patterns [25–27]. Neurons that project to subcortical action-selection and motor structures are likely to carry different information than those targeting associative cortical areas involved in higher-order function. Consequently, elucidating ACtx’s contribution to auditory-guided behavior necessitates studying its major projection classes as separate populations. Cortical output is dominated by two broad classes of excitatory projection neurons: intratelencephalic (IT) and extratelencephalic (ET). In ACtx, ET neurons are concentrated in layer (L) 5b and send dense projections to the midbrain, thalamus, striatum, and lateral amygdala, whereas IT neurons span L2/3-L6 and project widely to ipsiand contralateral cortex as well as striatum [28–32]. Although both IT and ET neurons are present in L5, the canonical cortical output layer, their largely non-overlapping projection patterns suggest distinct roles during behavior. Indeed, this projection rationale has strong anatomical and functional support across multiple sensory and motor cortices [26, 28, 33–37], underscoring the need for a cell-type–specific dissection of ACtx output channels during categorization.

Multiple lines of evidence indicate that ACtx L5 neurons play causal roles in auditory-guided behaviors. L5 ET projections to the midbrain and thalamus can rapidly modulate sensory tuning and gain [33–36, 38–43], whereas IT projections to frontal and association cortices shape the representation of auditory features and choice-related signals [44, 45]. Selective manipulation of corticostriatal projections, comprising both IT and ET neurons, disrupts task performance [37, 46, 47], and inhibiting either L5 IT or ET neurons during auditory learning slows task acquisition [48]. Research outside of ACtx further suggests that projection class strongly constrains function during behavior [49–57]. Despite extensive evidence for category-like responses in ACtx, powerful corticofugal influences on subcortical processing, and projection-specific roles within behavior, prior studies have rarely parsed the major L5 output channels during auditory categorization itself. In particular, the respective contributions of IT versus ET neurons to forming categorical representations, committing to a choice, and initiating action remain unresolved [33, 37, 47]. These considerations motivate a cell-type–specific test of whether and how ACtx L5 IT and ET neurons differentially route category information to decision and motor circuits.

To delineate the contributions of major cortical projection types to auditory category learning, we trained mice on a rate-categorization task and performed longitudinal two-photon calcium imaging of L2/3, L5 IT, and L5 ET neurons throughout learning. Across learning, L5 ET neurons, but not L2/3 or L5 IT neurons, exhibited dramatic tuning shifts, developing strong selectivity for the trained perceptual categories. This categorical selectivity persisted after excluding neurons whose activity was better explained by choice-related variables, indicating genuine stimulus-based encoding. Moreover, on perceptually ambiguous boundary trials, L5 ET population activity encoded choice, and the neural state immediately preceding sound onset could predict future choices. Together, these findings implicate the ACtx L5 ET projection system as a specialized cortical output channel that links sensory categorization to choice.

## Results

### Mice readily learn a head-fixed auditory categorization task

To examine learning-related changes across distinct ACtx cell types, we trained head-fixed mice to categorize the temporal rates of sinusoidally amplitude-modulated (sAM) white noise as “slow” or “fast” by licking left or right [48] (**Fig. 1a**). Mice reported choices within a brief window following sound onset: correct responses resulted in a small water reward while incorrect choices were punished with a five second timeout before the next trial (**Fig. 1b**). Category/spout mappings (left/right) were counterbalanced across mice to avoid systematic side biases. Mice were first trained on the easy stimuli (2, 2.8, 22.6, and 32 Hz) until they reached a proficiency threshold of 85% correct, which took mice, on average, 12 sessions (**Fig. 1c, Fig. S1a**). After this initial training, we introduced all nine stimuli (2-32 Hz in 0.5 octave steps) and tracked performance across days, grouped into early, middle, and late training epochs (**Fig. 1c**). Behavioral accuracy on nonboundary trials increased from early to late learning, driven by an improvement in categorizing both easy and hard stimuli (**Fig. 1d**). Session-wise psychometric fits showed steeper slopes and reduced bias across learning (**Fig. 1e,f**), whereas choice bias and lapse rates remained stable (**Fig. S1b,c**). Together, these results indicate the mice can readily learn this task and continue to improve their ability to categorize stimuli across multiple days of training.

**Figure 1:**
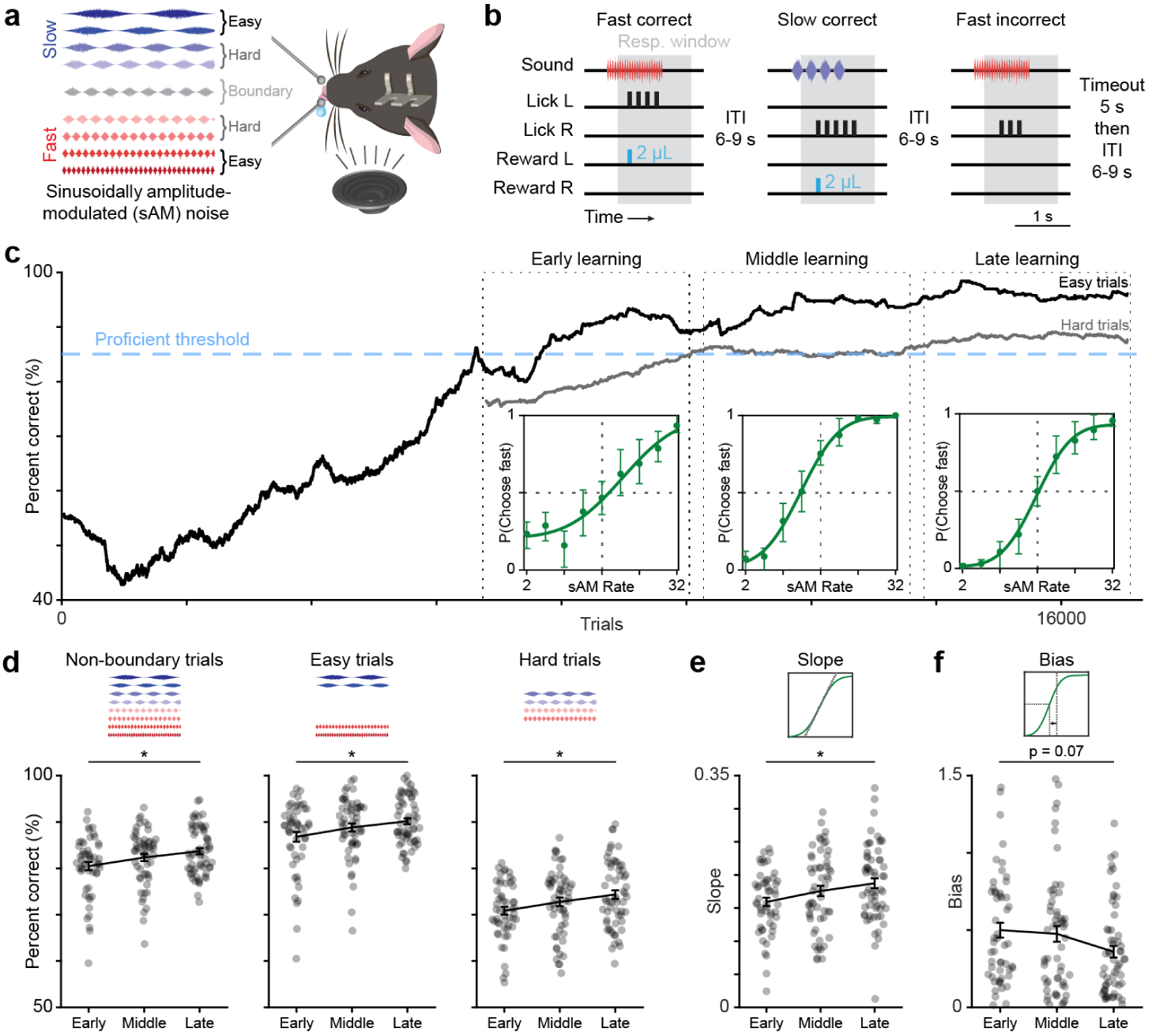
Mice readily learn a head-fixed auditory categorization task. (a) Schematic of head-fixed categorization behavior. (b) Schematic of trial structure and outcomes. (c) Example learning trajectory for easy and hard trials in a single mouse. Insets: psychometric curves from example sessions within each learning phase. Error bars represent binomial proportion confidence intervals. (d) Percent correct for non-boundary trials (one-way ANOVA, main effect for learning, *p* = 0.02) easy trials (one-way ANOVA, main effect for learning, *p* = 0.02), and hard trials (one-way ANOVA, main effect for learning, *p* = 0.03) for all sessions and mice across phases of learning (early: *n* = 53 sessions, middle: *n* = 57 sessions, late: *n* = 60 sessions; *N* = 11 mice). Asterisks represent main effects. (e) Slope of psychometric curves fit to each behavioral session broken up by learning phase (one-way ANOVA, main effect for learning, *p* = 0.03). Asterisk represents main effect. (f) Same as (e) for psychometric bias (one-way ANOVA, main effect for learning, *p* = 0.07).

### Category learning reshapes L5 ET tuning during active task engagement

We used our categorization task to investigate how distinct ACtx neuronal populations change with category learning. We selectively expressed GCaMP8s in L2/3, L5 IT, and L5 ET neurons using combinatorial viral, tracing, and genetic tools (**Fig. 2a**, see Methods). We then performed *in vivo* two-photon imaging of somatic calcium signals from these three populations in awake, head-fixed mice (**Fig. 2b**). After the initial training period (once mice reached our proficient threshold), we recorded activity during behavioral engagement on each subsequent day of categorical learning (Active, **Fig. 2c, d**). On the same days, and while imaging the same neurons, we also presented identical sounds after retracting the lick spouts (Passive), and we alternated the starting context (active versus passive) across days to control for changes in motivation and satiety (**Fig. 2c,d**). As an additional control, we imaged untrained (Naive) mice to assess baseline sAM tuning properties within the three distinct neural populations (**Fig. 2d**).

**Figure 2:**
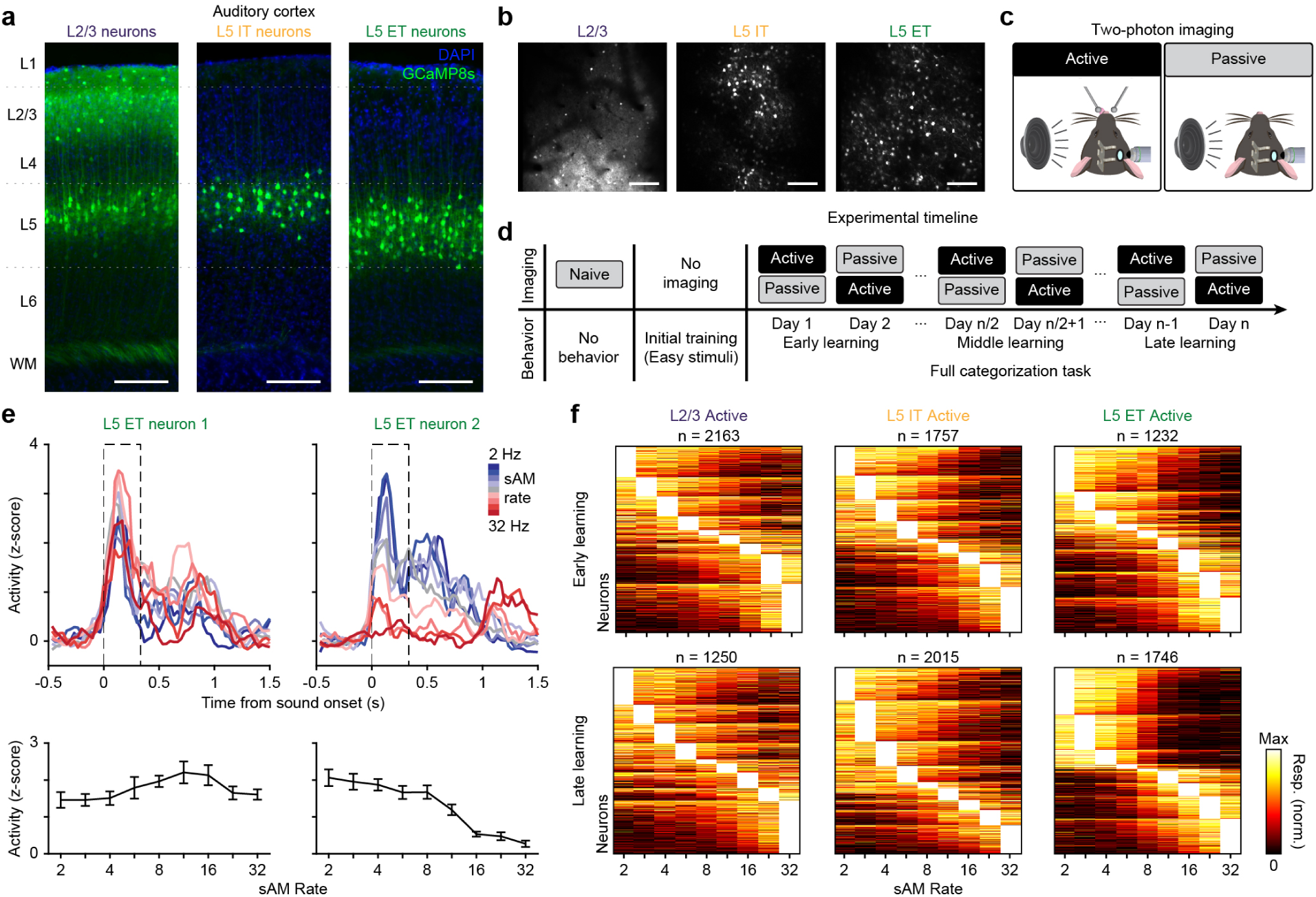
Category learning reshapes L5 ET tuning during active task engagement. (a) Coronal sections of ACtx showing GCaMP8s expression in L2/3, L5 IT, and L5 ET neurons. Scale bars represent 150*µm*. (b) Example two-photon imaging planes showing GCaMP8s expression in the three cell types. Scale bars represent 150*µm*. (c) Schematic of active and passive imaging sessions. (d) Schematic of imaging and behavioral training timeline. (e) Top: Example trial-averaged sound responses from two L5 ET neurons. Dashed box denotes the onset response window (0.33 s). Bottom: sAM rate tuning curves from the same two L5 ET neurons calculated by averaging activity within the onset response window. (f) Maximum-normalized response tuning curves sorted by preferred sAM rate for the three cell types comparing active early learning and active late leaning.

Across sessions, neurons exhibited heterogeneous evoked response profiles (onset, sustained, and offset; **Fig. S2a,b**), but we restricted subsequent analyses to neurons that were significantly responsive in a window immediately after sound onset, which captured the peak neural activity and decision-making period (**Fig. 2e, Fig. S1d**). The fraction of onset-responsive neurons varied across cell types and behavioral contexts (**Fig. S2c**). We then generated tuning curves for each neuron by averaging activity within this onset window to assess how stimulus selectivity evolved with learning (**Fig. 2e**). Visualizing normalized tuning curves suggested a learning-dependent change in L5 ET neurons from early to late training: responses became more broadly tuned to stimuli within the slow or fast categories (**Fig. 2f**). Further quantification confirmed that only L5 ET neurons developed robust category-specific tuning, preferentially responding to the four members of one category, whereas L2/3 and L5 IT neurons did not (**Fig. S2d,e**). Notably, this categorical organization emerged only during active task engagement and was absent during passive listening and in naive mice (**Fig. S2e**). These results indicate that L5 ET neurons adjust their sensory tuning in a context-dependent manner to generalize sounds within learned categories during behavior.

### Categorical coding emerges selectively in L5 ET neurons

Our tuning curve analyses indicated that L5 ET neurons increasingly assigned stimuli to a learned auditory category. To quantify this category preference and its evolution with learning, we computed a categorical selectivity index (CSI) for each neuron, where values near -1 or +1 indicate a strong preference for the slow or fast category, respectively, and values near 0 indicate no categorical bias (**Fig. 3a,b**). Examining CSI distributions across learning revealed minimal change for L2/3 and L5 IT neurons, whereas L5 ET neurons developed a pronounced bimodal distribution late in learning (**Fig. 3c, Fig. S3a**). During passive presentation of the same sounds, categorical selectivity was reduced for all cell types (**Fig. 3d, Fig. S3b**), and untrained mice showed no evidence of category preference (**Fig. S3c**). Thus, across learning, only the L5 ET population dynamically reshaped its responses to encode behaviorally relevant categories, and this organization emerged specifically during active task engagement.

**Figure 3:**
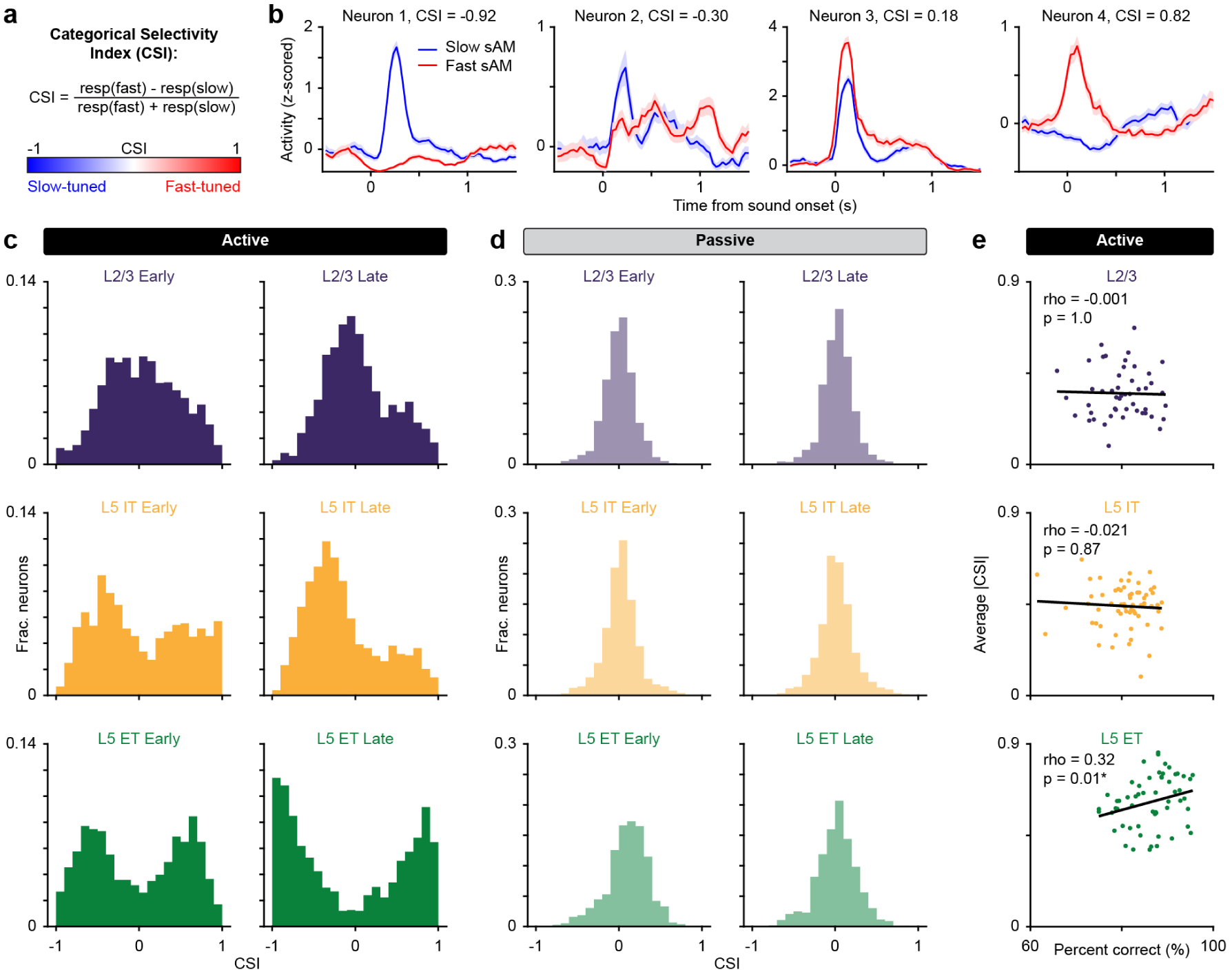
Categorical coding emerges selectively in L5 ET neurons. (a) Schematic of the categorical selectivity index (CSI). (b) Example neuron responses on correct slow and fast trials and their corresponding CSI values. (c) Histograms of CSI values for onset responsive neurons during active behavior. Plots are separated by early and late learning sessions across cell types. (d) Same as (c) for passive listening sessions. (e) Correlation between average |CSI| and performance (percent correct) on each session. Lines represent linear regression fits (L2/3: *n* = 49 sessions, L5 IT: *n* = 63, L5 ET: *n* = 58; Spearman’s Rho). reflect learned perceptual categories specifically during active task engagement.

For each imaging session, we quantified population-level categorical selectivity as the session-average absolute CSI (|CSI|). Categorical selectivity was highest for L5 ET neurons and increased from early to late learning (**Fig. S3d**). A complementary analysis, fitting a two-component Gaussian mixture to the CSI distribution, also revealed increasing bimodality in L5 ET across learning (**Fig. S3e**). Session-wise categorical selectivity in L5 ET correlated with behavioral performance (**Fig. 3e**, **Fig. S3f**). For comparison, L2/3 and L5 IT neurons also showed higher categorical selectivity during active versus passive conditions, but their |CSI| values did not change with learning and did not track behavioral performance. Motivated by evidence that similarly-tuned neurons form local cortical microcircuits [58–60], we asked whether categorically-selective L5 ET neurons were spatially clustered. Slow-preferring L5 ET neurons were significantly closer to each other than expected by chance, whereas fast-preferring neurons did not show reliable clustering (**Fig. S3g**). Taken together, only the L5 ET population developed categorical selectivity across learning.

Categorical selectivity within the L5 ET population could, in principle, arise via two mechanisms: 1) differential amplification and suppression of pre-existing preferences, such that strongly tuned neurons become more responsive while weakly tuned neurons drop out of the effective coding pool; or 2) within-cell retuning, whereby individual neurons shift their stimulus preferences to align with the learned categories (**Fig. S4a**). The absence of categorically-selective neurons in naive mice (**Fig. S3c**), together with learning-dependent changes in L5 ET tuning curves (**Fig. 2f**, **Fig. S2d,e**), is consistent with the second mechanism. To test this directly, we longitudinally tracked the same L5 ET neurons across all days of training and quantified changes in their tuning and CSI (**Fig. S4b**). Individual L5 ET neurons altered their stimulus preferences to encode the slow and fast categories (**Fig. S4c**), and |CSI| increased across learning (**Fig. S4d**). These effects were specific to active task engagement, were not observed during passive listening, and were absent in longitudinally matched L2/3 and L5 IT neurons(**Fig. S4d**). These results show that, unlike L2/3 and L5 IT neurons, individual L5 ET neurons dynamically change their activity across learning to

### Functional diversity of L5 ET neurons underlies discrimination and categorization

Although most L5 ET neurons develop category-selective responses across learning, mice first learn to discriminate the “easy” slow and fast exemplars (far from the category boundary) before full categorical learning begins (**Fig. 1c**). This implies the presence of neurons that reliably distinguish slow from fast early in learning. We asked whether these early “discriminator” neurons later acquire categorical selectivity (adapting-population hypothesis), or whether a largely separate set of neurons becomes category-selective as learning progresses (separate-population hypothesis, **Fig. 4a**). A key indication of the latter hypothesis would be a subset of neurons that consistently support slow-fast discrimination across all learning phases.

**Figure 4:**
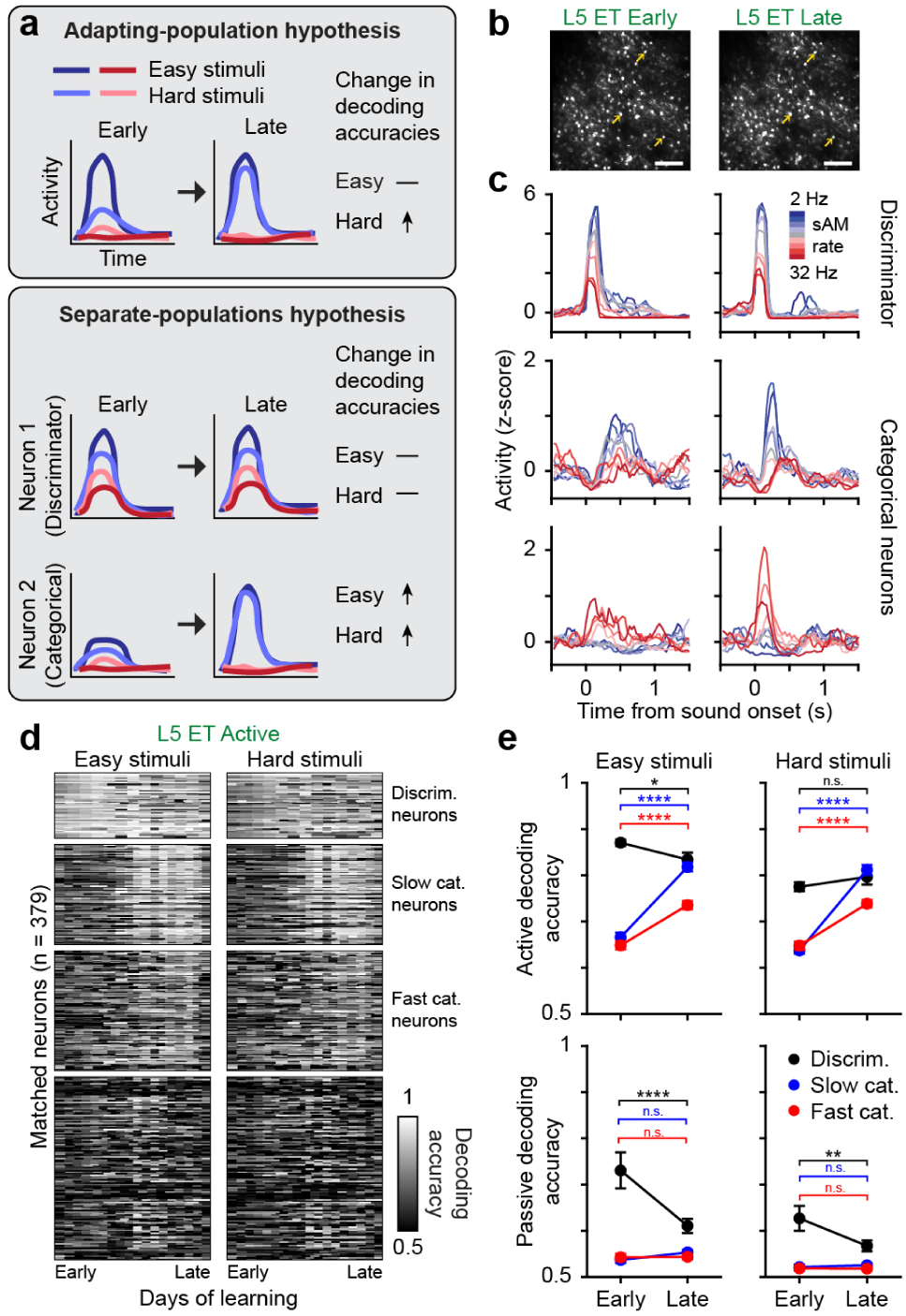
Functional diversity of L5 ET neurons underlies discrimination and categorization. (a) Schematic of two hypotheses for how neurons adapt to encode the hard sAM stimuli. (b) Example L5 ET field of view matched between early and late in learning. Arrows show examples in (c) matched across FOVs. Scale bars represent 150*µm*. (c) Example L5 ET neuron responses in an early and late session. (d) Decoding accuracies for matched L5 ET neurons across learning on both easy and hard stimuli. Neurons are broken up into a discriminator population, categorical populations (slow and fast), and all other neurons. (e) Average easy and hard category decoding accuracies for discriminator neurons (black, *n* = 53), slow categorical neurons (blue, *n* = 81), and fast categorical neurons (red, *n* = 97) in active (top) and passive (bottom) across learning (Active easy: repeated measures mixed effects model with post-hoc Tukey test, main effect for neural group, *p <* 0.0001, main effect for learning, *p <* 0.0001, interaction, *p <* 0.0001; Active hard: repeated measures mixed effects model with post-hoc Tukey test, main effect for neural group, *p <* 0.0001, main effect for learning, *p <* 0.0001, interaction, *p <* 0.0001; Passive easy: repeated measures mixed effects model with post-hoc Tukey test, main effect for neural group, *p <* 0.0001, main effect for learning, *p* = 0.0007, interaction, *p <* 0.0001; Passive hard: repeated measures mixed effects model with post-hoc Tukey test, main effect for neural group, *p <* 0.0001, main effect for learning, *p* = 0.009, interaction, *p* = 0.006; For all sub-panels, asterisks represent pairwise comparisons and color denotes the pair).

We again analyzed neurons matched longitudinally across learning to address this question (**Fig. 4b**). We identified a subset of neurons that encoded sAM rate with high fidelity in both early and late sessions (**Fig. 4c**, top), potentially comprising a stable pool of “discriminator” neurons. We also observed neurons with initially weak stimulus preferences that, over learning, developed strong categorical tuning (**Fig. 4c**, middle and bottom), suggesting that category-selective neurons do not generally begin as easy discriminators and supporting the separate-populations hypothesis. We therefore set out to identify neurons that maintained strong slow-fast discrimination across learning, an analysis that would test whether the data are better explained by an adapting population or by largely separate functional populations.

To test for a subset of stable L5 ET discriminator neurons, we performed single-neuron binary decoding of slow versus fast categories, computed separately using only the easy (far from the category boundary) or hard (near the category boundary) stimuli (**Fig. S5a**). If such neurons exist, they should maintain stable easy-stimuli decoding accuracies across learning. Indeed, while many L5 ET neurons increased their decoding accuracy with learning, we identified a small subset of neurons (see Methods) that already decoded easy stimuli with high accuracy in early sessions and remained high late in learning, with only a modest decline in performance (**Fig. 4d,e, Fig. S5**). In these discriminator neurons, decoding of the hard stimuli did not increase across learning, which is inconsistent with a pure adapting-population account in which early discriminators broaden to encode full categories. In contrast, neurons that were strongly categorical by late learning (high |CSI|, see Methods) began with weaker easy-stimulus decoding which improved across days (**Fig. 4e**, top), consistent with the emergence of a distinct categorical pool. During passive listening, discriminator neurons still decoded above chance (**Fig. 4e**, bottom, **Fig. S5c**), suggesting that they encode stimulus identity robustly across behavioral contexts, whereas categorical neurons performed near chance without behavioral engagement, reinforcing the context dependence of categorical coding. Together, these results indicate that the L5 ET population comprises at least two functional subgroups, stable discriminators and emergent categorical neurons, highlighting the diversity of behaviorally relevant responses in ACtx L5 ET neurons.

### ACtx encodes both stimulus and choice

On correct, categorizable trials, the stimulus category (slow versus fast) is perfectly coupled to choice (left versus right lick), creating a potential confound between stimulus- and choice-related activity. Prior studies have reported robust choice encoding during auditory discrimination in ACtx, and more generally, widespread modulation by movement and internal state [18, 44, 61–72].

To characterize choice (lick-related) signals in our data, we first examined L5 ET activity around spontaneous licking in trained mice during silence and observed modulation that preceded lick onset (**Fig. S6a–c**). To test whether this effect depends on learning, we imaged a separate cohort performing a “licking-only” paradigm without sounds and again observed lick-locked modulation in ACtx L5 ET neurons (**Fig. S6d–f**), consistent with prior reports [62, 69]. These observations indicate that choice/movement signals are present in ACtx and must be explicitly examined when assessing category selectivity.

We next separated neurons according to whether their activity was better explained by stimulus-related or choice-related variables using a generalized linear model (GLM) [73–76]. The design matrix included time-lagged predictors for stimulus identity and lick direction, and we fit the model to z-scored deconvolved calcium activity (**Fig. 5a**, **S7a**; see Methods). After fitting a model for each responsive neuron, we quantified the coefficient contributions (*ω*) of stimulus and lick predictors (**Fig. 5b, S7b**). Additionally, we compared cross-validated performance of partial models containing only stimulus predictors or only licking predictors (**Fig S7c,d**). Neurons were labeled “stimulus neurons” if both the stimulus-only model outperformed the licking-only model and *ω_stim_ > ω_lick_*; conversely, neurons were labeled “choice neurons” when the licking-only model dominated and *ω_stim_ < ω_lick_* (**Fig. 5b, S7b-e**). All three cell types contained both stimulus and choice neurons, and the proportions of these classes were stable across learning (**Fig. S8a**). This emphasizes that ACtx populations multiplex sensory- and choice-related signals as mice perform the categorization task.

**Figure 5:**
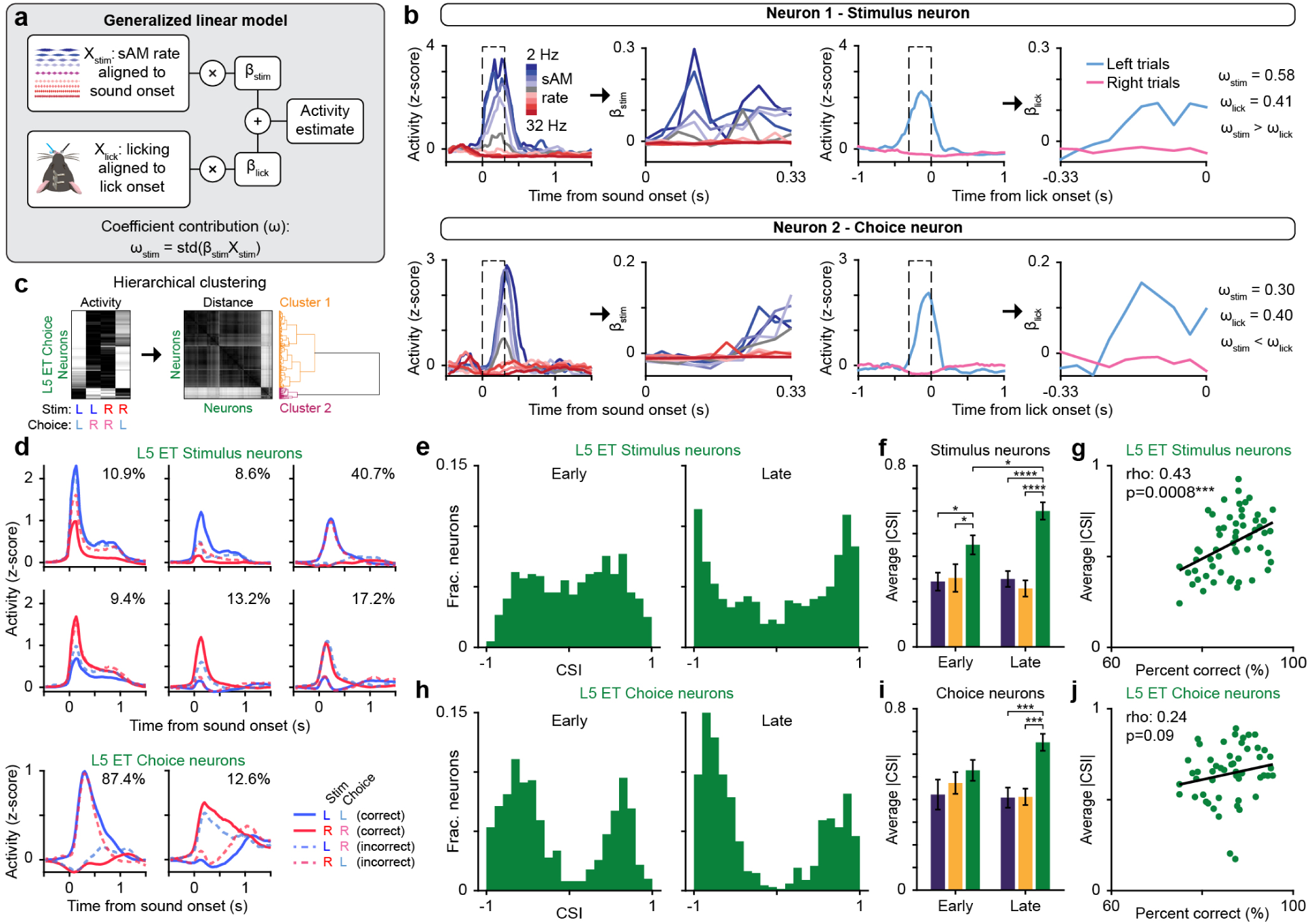
Categorical selectivity develops within L5 ET stimulus neurons, not choice neurons. (a) Schematic of a generalized linear model fit with stimulus and licking activity. (b) Examples of two neurons fit by the GLM and classified as either a stimulus neuron or choice neuron. (c) Hierarchical clustering of L5 ET choice neurons. Left: average rescaled activity for correct and incorrect left and right choices for all neurons sorted by cluster identity. Middle: matrix of pairwise distances between all neural responses sorted by cluster identity. Right: clustering tree (dendrogram) computed from the pairwise distances with colors denoting each cluster group. (d) Average cluster responses of L5 ET stimulus neurons (top) and L5 ET choice neurons (bottom). Percentages represent the size of each cluster relative to the total number of stimulus (or choice) neurons. (e) Histograms of CSI values for L5 ET stimulus neurons responsive during active behavior. Plots are separated by early and late sessions. Only correct trials were used to calculate CSI. (f) Average |CSI| for stimulus neurons in each session separated by cell type an learning phase (L2/3 early: *n* = 15 sessions, L5 IT early: *n* = 16, L5 ET early: *n* = 18, L2/3 late: *n* = 17, L5 IT late: *n* = 19, L5 ET late: *n* = 20; Two-way ANOVA with post-hoc Tukey test, main effect for cell type, *p <* 0.0001, interaction, *p* = 0.06, asterisks represent pairwise comparisons). (g) Correlation between average |CSI| and performance (percent correct) on each session in L5 ET stimulus neurons. Line represents linear regression fit (*n* = 58 sessions; Spearman’s Rho). (h) Same as (e) for L5 ET choice neurons. (i) Same as (f) for choice neurons (L2/3 early: *n* = 13 sessions, L5 IT early: *n* = 20, L5 ET early: *n* = 16, L2/3 late: *n* = 15, L5 IT late: *n* = 22, L5 ET late: *n* = 18; Two-way ANOVA with post-hoc Tukey test, main effect for cell type, *p* = 0.0004, asterisks represent pairwise comparisons). (j) Same as (g) for L5 ET choice neurons (*n* = 52 sessions; Spearman’s Rho).

We validated the GLM-based classification by leveraging correct and incorrect trials. True choice neurons should coincide with the mouse’s decision (left versus right) regardless of correctness. Consistent with this prediction, unsupervised clustering of trial-averaged responses (separating correct and incorrect trials) revealed two clear groups of choice neurons corresponding to left- and right-directed decisions (**Fig. 5c,d, S8b**). We also observed an asymmetry where more neurons encoded left choices, suggesting a lateralization of choice encoding (see Discussion). In contrast, stimulus neurons exhibited heterogeneous response properties (**Fig. 5d, S8c**). Some stimulus neurons encoded category identity irrespective of behavioral outcome, others were outcome-sensitive (e.g., stronger on correct trials), and some showed residual choice-like modulation despite being better explained by stimulus predictors in the GLM. Together, these analyses support two functionally-defined subpopulations during behavior: choice neurons, which robustly encode the mouse’s left/right decision, and stimulus neurons, which carry diverse sensory-related signals.

### Categorical selectivity develops within L5 ET stimulus neurons, not choice neurons

We next asked whether the learning-related increase in categorical selectivity observed in L5 ET neurons reflects genuine sensory encoding or a by-product of choice-related activity. Having classified neurons on a per-session basis as either stimulus or choice neurons with the GLM, we recomputed CSI within each subgroup. Restricting the analysis to stimulus neurons revealed that L5 ET showed an even more pronounced increase in categorical selectivity across learning, and session-averaged |CSI| closely tracked behavioral performance (**Fig. 5e,g**). As before, categorical selectivity was markedly reduced during passive listening, and neither L2/3 nor L5 IT stimulus neurons exhibited learning-related changes (**Fig. 5f, S9a-c**). In contrast, choice neurons in all three populations showed no systematic change in categorical selectivity and did not correlate with behavior (**Fig. 5h-j, S9d-f**). These results indicate that the emergence of categorical coding in ACtx arises specifically within the stimulus-driven subset of L5 ET neurons, and is not a by-product

Given the distinct response profiles of stimulus and choice neurons, we next asked how each group represents category identity and behavioral choice over time. Based on our clustering results, choice neurons should primarily encode choice and reflect category only on correct trials, whereas stimulus neurons, which showed heterogeneous responses, might encode information about both variables. We therefore trained a linear classifier to decode category or choice from population activity using all trials (correct and incorrect; **Fig. 6, S10**). Stimulus neurons supported robust decoding of both variables, with distinct temporal profiles: category information rose shortly after sound onset, while choice information ramped toward the decision (**Fig. 6a, S10a**). L5 ET populations yielded the highest decoding accuracies and strongest correlations to performance relative to L2/3 and L5 IT populations (**Fig. 6b,c, S11a**). Category and choice were decoded with comparable fidelity from L2/3 and L5 IT stimulus neurons, but L5 ET stimulus neurons showed a modest, reliable bias towards choice decoding (**Fig 6c**). These findings indicate that stimulus-driven populations jointly encode both category and choice, with L5 ET stimulus neurons particularly well positioned to transform sensory information into a decision signal (see Discussion).

**Figure 6:**
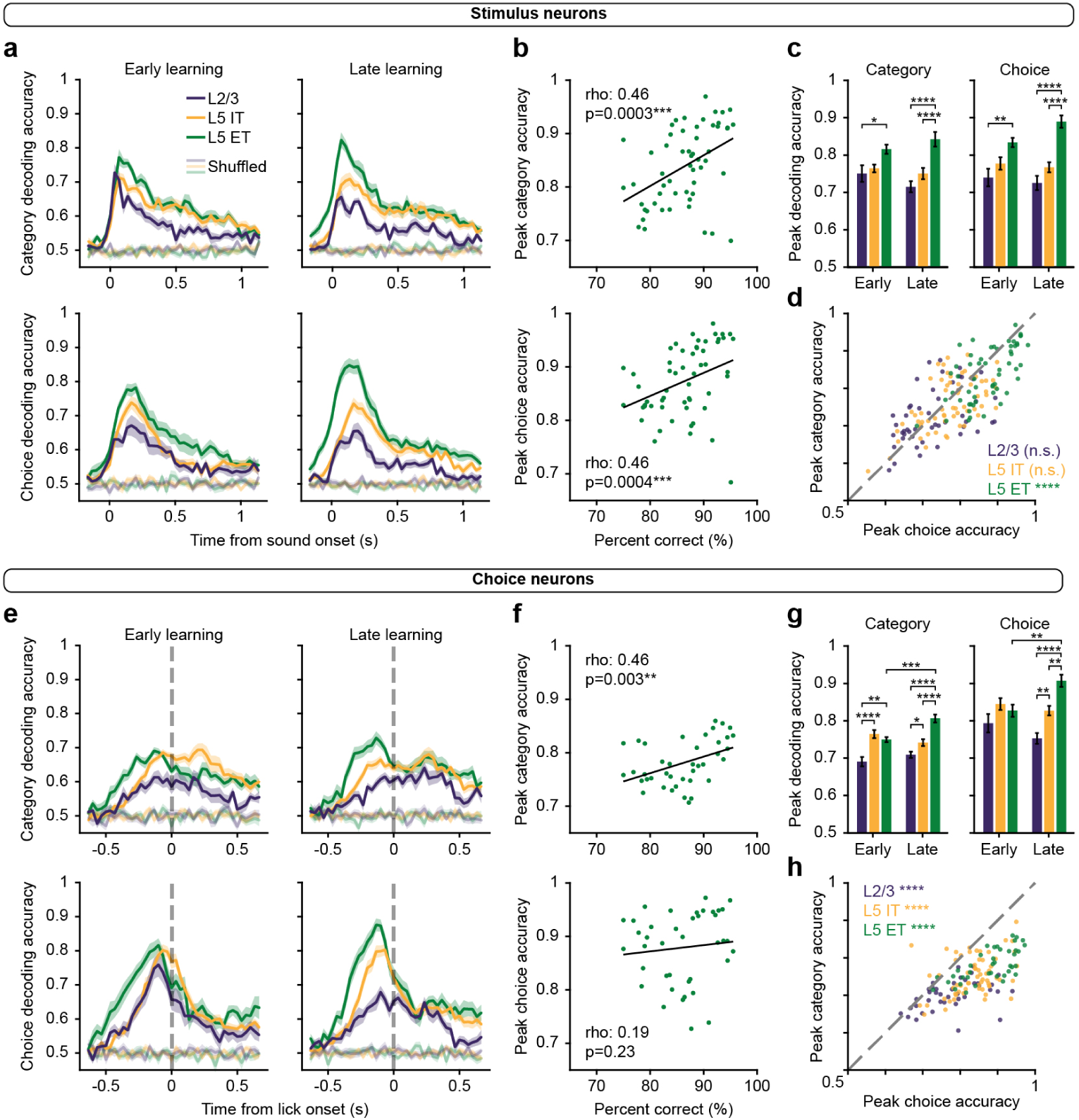
Stimulus neurons encode both category and choice. (a) Average category (top) and choice (bottom) decoding accuracies for stimulus neurons across the duration of a trial. Transparent lines represent shuffled data. (b) Correlation between performance and decoding accuracies for category (top) and choice (bottom) of L5 ET stimulus neurons. Lines represent linear regression fit (*n* = 56 sessions; Spearman’s Rho). (c) Peak decoding accuracies for stimulus neurons across learning for both category and choice decoding (L2/3 early: *n* = 15 sessions, L5 IT early: *n* = 20, L5 ET early: *n* = 17, L2/3 late: *n* = 19, L5 IT late: *n* = 23, L5 ET late: *n* = 21; Category: Two-way ANOVA with post-hoc Tukey test, main effect for cell type, *p <* 0.0001; Choice: Two-way ANOVA with post-hoc Tukey test, main effect for cell type, *p <* 0.0001, asterisks represent pairwise comparisons). (d) Comparison of peak decoding accuracy between category and choice on each session in stimulus neurons (L2/3: *n* = 49 sessions, paired t-test, *p* = 0.84; L5 IT: *n* = 63, paired t-test, *p* = 0.13; L5 ET: *n* = 56, paired t-test, *p <* 0.0001). (e) Same as (a) for choice neurons. (f) Same as (b) for L5 ET choice neurons (*n* = 40 sessions; Spearman’s Rho). (g) Same as (c) for choice neurons (L2/3 early: *n* = 13 sessions, L5 IT early: *n* = 20, L5 ET early: *n* = 13, L2/3 late: *n* = 17, L5 IT late: *n* = 23, L5 ET late: *n* = 13; Category: Two-way ANOVA with post-hoc Tukey test, main effect for cell type, *p <* 0.0001, main effect for learning, *p* = 0.04, interaction *p* = 0.0006; Choice: Two-way ANOVA with post-hoc Tukey test, main effect for cell type, *p <* 0.0001, interaction, *p* = 0.002, asterisks represent pairwise comparisons). (h) Same as (c) for choice neurons (L2/3: *n* = 40 sessions, paired t-test, *p <* 0.0001; L5 IT: *n* = 62, paired t-test, *p <* 0.0001; L5 ET: *n* = 40, paired t-test, *p <* 0.0001).

Choice neurons robustly encoded behavioral choice: choice-decoder accuracy ramped before the first lick and peaked around the decision (**Fig. 6e**). By contrast, category decoding from choice neurons was weaker and temporally diffuse, without a clear peak relative to sound or lick onsets (**Fig. 6e, S10b**). Although this category decoding increased across learning and correlated with behavioral performance (**Fig. 6f,g, S11b**), this pattern is consistent with improving alignment between category and choice as training progresses rather than with the emergence of genuine category signals in choice neurons. Supporting this interpretation, choice-decoder accuracy did not correlate with performance (**Fig. 6f, S11b**). Notably, for both category and choice decoders, L5 populations (both IT and ET) outperformed L2/3 (**Fig. 6g**), consistent with a greater role for deep layers in guiding future actions. Across all three cell types, choice neurons did not encode category identity nearly as strongly as choice (**Fig. 6h**), confirming that they primarily participate in decision planning/execution rather than representing stimulus categories.

### ACtx encodes choice history under sensory ambiguity

Our decoding results show that L5 ET populations encode choices more strongly than categories, suggesting that they are involved in using stimuli to inform decisions. However, this is difficult to assess given the strong mapping between category and choice. To directly investigate this transformation, we included an ambiguous boundary stimulus (8 Hz) that carried no categorical evidence and was pseudo-randomly paired with left or right reward, prompting mice to make different choices to the same sound. On these boundary trials, any left-right neural separation reflects choice-related activity rather than stimulus category. State-space visualizations with principal component analysis (PCA) revealed that L5 ET population trajectories diverged for left versus right choices despite identical sensory input, with divergence peaking around the decision (**Fig. 7a**). To quantify this effect, we calculated the euclidean distance between the average lick-aligned left and right choice trajectories and found reliable choice separation in all three cell types, with the strongest effect in L5 ET (**Fig. 7b,c**). The separation was evident in both stimulus- and choice-classified subsets, although larger for the latter (**Fig. S12a–d**). Thus, L5 ET populations carry robust choice signals even when sensory evidence is held constant.

**Figure 7:**
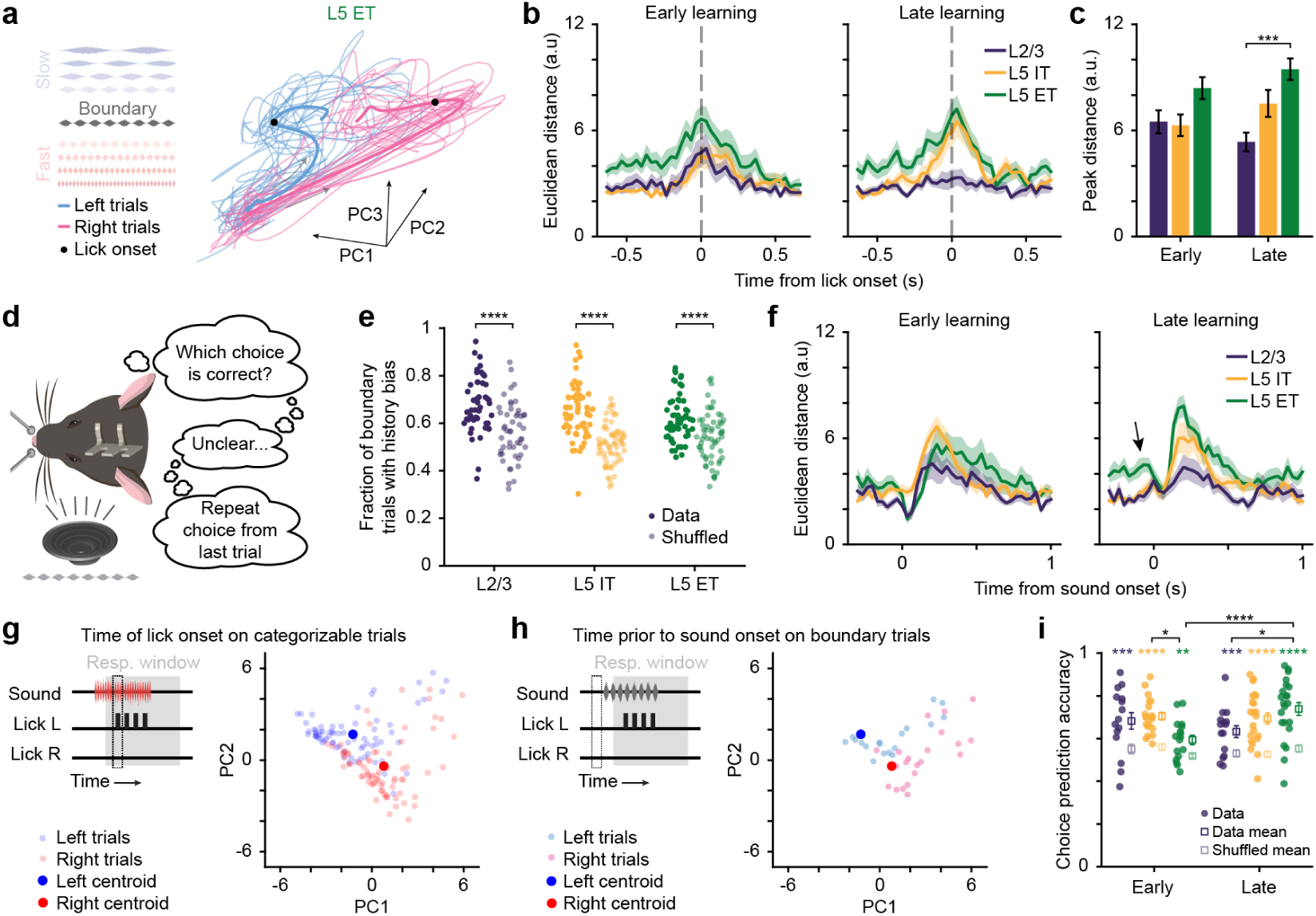
ACtx encodes choice history under sensory ambiguity. (a) Neural trajectories of left and right boundary trials of an example L5 ET session aligned to lick onset. Transparent lines represent individual trials and bold lines represent average trajectories. (b) Euclidean distance between average lick-aligned left and right boundary trial trajectories across learning and cell types. (c) Peak distance between trajectories across all sessions (L2/3 early: *n* = 15 sessions, L5 IT early: *n* = 20, L5 ET early: *n* = 17, L2/3 late: *n* = 15, L5 IT late: *n* = 23, L5 ET late: *n* = 22; Two-way ANOVA with post-hoc Tukey test, main effect for cell type, *p <* 0.0001). (d) Illustration of how mice may make decisions on boundary trials. (e) Fraction of boundary trials that have the same choice as the previous trial compared to a shuffled distribution of boundary choices (L2/3: *n* = 49 sessions, paired t-test, *p <* 0.0001; L5 IT *n* = 63, paired t-test, *p <* 0.0001; L5 ET *n* = 58, paired t-test, *p <* 0.0001). (f) Same as (b) for stimulus-aligned trajectories. Black arrow emphasizes pre-stimulus choice information in L5 ET neurons. (g) Population activity at lick onset for all correct slow/left (blue) and fast/right (red) trials for an example session. Large circles denote the centroids calculated from the mean of all points. (h) The same example session and centroids from (f) plotted with the population activity on boundary trials immediately preceding stimulus onset. Each point is colored by the future choice of the animal. (i) Accuracy of predicting a boundary choice based on the pre-stimulus population activity (L2/3 early: *n* = 15 sessions, paired t-test, *p* = 0.0007; L5 IT early: *n* = 20, paired t-test, *p <* 0.0001; L5 ET early: *n* = 17, paired t-test, *p* = 0.004; L2/3 late: *n* = 15, paired t-test, *p* = 0.0005; L5 IT late: *n* = 23, paired t-test, *p <* 0.0001; L5 ET late: *n* = 22, paired t-test, *p <* 0.0001, colored asterisks represent comparisons against shuffled data; Two-way ANOVA with post-hoc Tukey test, interaction, *p* = 0.002, black asterisks represent pairwise comparisons of movement or choice signals.

We next investigated how mice made decisions on these ambiguous trials. Prior work shows that recent actions can bias subsequent decisions toward repetition, especially in the presence of uncertain stimuli [17, 48, 77–82] (**Fig. 7d**). Consistent with this, mice were more likely to repeat the previous trial’s choice on boundary trials, leading to upcoming choice predictions above chance (**Fig. 7e**). This suggests that when sensory evidence is uninformative, an internal representation of the prior decision biases the next. Although choice history signals are well established in higher-order cortices [17, 77, 81, 83–86], they may also be present in ACtx. Supporting this possibility, stimulus-aligned analyses showed that L5 ET populations carried choice information prior to sound onset (**Fig. 6a, 7f, S11b, S12e,f**). Because trial onset was unpredictable and stimuli were pseudo-randomized, anticipatory preparation should be minimal; therefore, pre-stimulus choice-predictive activity in ACtx likely reflects an internal representation of the previous decision and/or top-down input conveying choice history.

To test whether ACtx carries choice history information, we asked if pre-stimulus population activity predicts choices on boundary trials. We first defined left- and right-choice neural subspaces by computing a low-dimensional population space (PCA) from activity at lick onset on correct, categorizable trials and then taking the mean (centroid) for each choice (**Fig. 7g**). These centroids provide canonical choice-linked population states. For each boundary trial, we compared the pre-stimulus population activity to the two centroids, predicted the upcoming choice based on whichever centroid was closer, and compared this to the true future choice (**Fig. 7h**). This simple nearest-centroid classifier yielded significantly above-chance prediction of left versus right choices across all three cell types (**Fig. 7i, S12g,h**), supporting the view that choice-related activity is present in ACtx before stimulus onset on ambiguous trials. Furthermore, this representation significantly increased across learning in L5 ET neurons (**Fig. 7i, S12g,h**), emphasizing the notion that this population becomes increasingly involved in transforming incoming information into behavioral decisions. These results suggest that a choice-history–dependent bias in the initial cortical state can tilt the population toward one of the choice-linked states when sensory evidence is uninformative.

## Discussion

Auditory categorization allows stimuli with different acoustic properties to be perceptually grouped together to guide behavior. Prior studies have implicated ACtx as a hub for categorization [8–14, 24], but these past studies have lacked the cell type specificity required to truly link structure to function. Indeed, theoretical work has suggested that distinct neuron classes within local circuits can differentially compute behavioral information, underscoring the need for cell-type-specific approaches [87, 88]. To address this gap in knowledge, we used *in vivo* two-photon calcium imaging to record the activity of distinct projection neuron types while mice learned an auditory categorization task. We found that L5 ET neurons develop categorical representations of the trained auditory stimuli (**Fig. 3**). Notably, this categorical encoding was context-dependent: it was present during active task engagement but absent during passive listening in the same neurons on the same day. This indicates that top-down or behavioral context is required for categorical coding, as opposed to being a purely stimulus-driven phenomenon. Furthermore, the emergence of categorical selectivity was a learned effect that developed across training and was absent in untrained mice. Using a GLM to factor out neurons encoding choice, we confirmed that the development of categorical selectivity in L5 ET neurons was driven by stimulus-specific encoding rather than a confound of licking behavior (**Fig. 5**). In contrast, neither L2/3 nor L5 IT neurons displayed strong categorical selectivity or notable changes in their response properties across learning. Together, our results demonstrate that ACtx L5 ET projection neurons play a critical and specific role in forming categorical representations during an auditory-guided behavior.

In addition to categorical responses, we investigated how these cell types encoded choice. Responses related to behavioral choice have been observed in multiple primary sensory cortices within both superficial and deep layers [44, 61–64, 66–68, 76, 89–91]. Consistent with this, we found robust choice encoding across all three cell types, with choice signals being most pronounced in L5 IT and L5 ET neurons (**Fig. 6**). To ensure that these choice signals were not simply a by-product of the correlation between stimulus category and choice (since mice directionally licked to report their decision), we examined neural activity at the category boundary where stimuli were perceptually ambiguous. Even when the same ambiguous sound was presented, ACtx activity differentiated left versus right lick choices, with L5 ET neurons showing the strongest representation of the upcoming choice (**Fig. 7**). In other words, ACtx (especially L5 ET cells) encoded the mouse’s decision independently of auditory category identity. We also found that mice tend to repeat their prior choice when presented with an ambiguous stimulus. This behavioral bias suggests that previous choices influence future decisions, a phenomenon often referred to as choice history bias. Frontal and parietal association areas have been implicated in mediating such choice history biases [17, 77, 81, 84–86, 92]. In particular, the posterior parietal cortex (PPC) has direct projections to ACtx and is known to influence auditory decisions and biases [17, 92, 93]. In line with this, we found that choice history influenced baseline (pre-stimulus) activity in ACtx: the neural activity immediately before an ambiguous sound could predict the ensuing choice on that trial with above-chance accuracy. This implies that areas such as PPC may convey choice history signals to ACtx, biasing ACtx activity and thereby biasing decisions when sensory information alone is insufficient to confidently categorize the stimulus. Taken together, our findings show that ACtx maintains distinct representations of category and choice, likely acting as an integrative hub where feedforward sensory evidence converges with feedback decision signals.

Our results suggest that L5 ET neurons play a more prominent role in auditory categorization than either L2/3 or L5 IT neurons. Several lines of evidence help explain this distinction. First, L5 ET neurons are thick-tufted pyramidal cells with apical dendrites that extend to layer 1, making them morphologically well-situated to integrate inputs across the cortical column [26, 31, 94–96]. Circuit mapping has shown that they receive synaptic inputs from multiple cortical areas as well as direct thalamic innervation [97–101], allowing them to combine bottom-up auditory information with top-down contextual signals. Second, L5 ET neurons project subcortically to decision- and reinforcement-related structures, including the inferior colliculus (IC) and striatum [102–112]. Notably, IC neurons in bats exhibit categorical responses to vocalizations [113], and corticocollicular feedback is required for learning-related plasticity in rodents [109, 110]. Thus, L5 ET outputs are positioned not just to convey category information but to shape subcortical circuits during learning. Third, L5 ET neurons possess intrinsic properties that enhance their output efficacy. They fire at higher rates than other excitatory neurons [28, 114–116], which could drive plasticity in subcortical targets. Together, these morphological, anatomical, and physiological features explain why L5 ET neurons, more than L2/3 or L5 IT, develop categorical representations and play a privileged role in shaping auditory-guided behavior.

Our cell-type–specific results also provide a mechanistic account of prior causal evidence linking cortical outputs to task performance [48]. Silencing ACtx altered decision strategies and impaired performance across learning, and inhibition of either L5 IT or L5 ET neurons significantly slowed learning while sparing expert performance [48]. The categorical code that emerged specifically in L5 ET populations offers a plausible mechanistic basis, where disruption of these neurons would impede the formation and subsequent dissemination of categorical representations to subcortical targets. In parallel, the choice encoding observed in L5 IT neurons likely propagates across cortical areas to facilitate the integration of task context, rule information, and history-dependent feedback necessary for associative learning. Thus, both projection pathways make distinct yet complementary contributions; perturbing either is sufficient to degrade learning, even if moment-to-moment performance in experts remains relatively preserved.

We observed substantial heterogeneity within each neuronal class, with functionally distinct subgroups defined by their encoding properties emerging even among neurons of the same type. In all three cell types, a subset of “stimulus neurons” responded selectively to the sounds themselves (with various tuning preferences) and carried information about stimulus identity or category. Another subset of “choice neurons” did not necessarily respond to the sound itself but showed activity aligned to decisions (e.g., ramping up activity when the mouse was about to lick left versus right). Notably, only the L5 ET stimulus neurons developed strong categorical selectivity across learning, reinforcing that the emergence of categorical tuning was a genuine sensory representation specific to this subgroup and cell type. The stimulus neurons in L2/3 and L5 IT, by contrast, often remained more tied to the physical features of sounds and did not show a clear grouping by category even after learning. Interestingly, we found that stimulus neurons (particularly in L5 ET) were able to decode both the category and the eventual choice with high fidelity from their activity. In fact, L5 ET stimulus neurons were better at decoding choice than category (see decoding analysis in **Fig. 6**), even though they were defined based on their sound responses. This, alongside their subcortical projections, suggests that L5 ET neurons might serve as a link in the sensorimotor chain, converting perceptual category information into a planned motor action.

In contrast to stimulus neurons, choice neurons did not exhibit changes in categorical selectivity with learning, nor did their activity correlate with behavioral performance. Instead, these neurons seemed to primarily reflect the decision process or motor plan itself, especially in L5 IT and ET (**Fig. 6**). Across all cell types, more choice neurons were tuned for the contralateral action (i.e., licking to the side opposite the imaged hemisphere) than for the ipsilateral action (**Fig. 5, Fig. S9**), suggesting that choice neurons may be preferentially involved in contralateral movements. This bias was especially pronounced in L5 ET neurons. Indeed, previous anatomical studies report that most ACtx L5 ET axons descend ipsilaterally, while a subset project bilaterally to brainstem targets [117, 118].

Despite the clear differences we observed between stimulus and choice neurons in our task, the origin of these functional subgroups remains unclear. One plausible explanation is that they correspond to differences in anatomical subcircuits even within the same genetic cell type. For example, recent work in motor cortex demonstrated that L5 ET neurons with different projection targets have distinct roles in movement control [119]. However, teasing apart such differences is challenging as many L5 ET neurons send collateral projections to multiple downstream targets [28, 37]. Another possibility is that stimulus and choice neurons differ in the inputs they receive. Notably, both stimulus and choice neurons in our data changed their activity between passive and active sessions, indicating that neither group is purely feedforward, but rather are modulated by behavioral context, likely via top-down inputs. This top-down modulation could originate from several different structures encoding task rules, expectations, or motor plans [17, 71, 120–124], and these regions might differentially innervate each functional subgroup. These potential differences in functional connectivity could explain how neurons within the same cell type develop distinct and specialized roles in behavior.

In addition to the L5 ET stimulus neurons that developed categorical tuning, we identified a small population of neurons in L5 ET that we termed “discriminator neurons” based on their high stimulus discriminability early in learning. Many of these discriminator neurons maintained a veridical encoding of stimulus identity across learning rather than switching to categorical tuning.

Unlike the categorical neurons, L5 ET discriminator neurons could decode stimulus identity even during passive listening sessions. This suggests that they may receive direct feedforward thalamocortical input [97, 99]. Indeed, thalamocortical synapses onto L5 neurons could drive such neurons regardless of behavioral context, explaining why their performance in discriminating stimuli did not improve with learning, and why some even showed decreased decoding accuracy as learning progressed. This decrease was more pronounced in passive listening, and could point to an effect of suppressed input following habituation from multiple days of exposure [125]. An alternative explanation could be that some discriminator neurons are “dropping out” of the population and being replaced with new neurons, which occurs in representational drift [126, 127].

While we used a GLM to account for licking, other movements also influence ACtx activity [69–72]. We did not explicitly model additional movement-related variables (e.g., facial or body movements) which could have provided better explanatory power. Accounting for these variables would likely not change our results, as uninstructed movements should not change systematically across learning or differ across our cell type groups. Additionally, our task design imposed a very short timing gap between sound onset and the required behavioral response. As a result, mice often made their decision while the sound was still playing (**Fig. S1d**), making it intrinsically difficult to temporally separate sensory responses from decision- or motor-related activity. We mitigated this using analyses to separate neurons based on whether their activity was better explained by stimulus or choice. Finally, our study focused on changes occurring from the point when the mice already understood the basic discrimination (i.e., after initial behavioral shaping on easy stimuli) through to expert performance on the fine categorization. We did not capture the earliest phase of learning when mice first learned to associate sounds with actions and rewards. Prior work has shown that animals can acquire knowledge of task contingencies quite rapidly (sometimes within a session), but the expression of that knowledge in behavior can be delayed or gradual [128, 129]. It is possible that L2/3 or L5 IT neurons play a crucial role in this rapid learning that we did not capture here.

Auditory categorization facilitates the rapid transduction of sensory stimuli into discrete behavioral decisions. Our findings show that L5 ET neurons play a key role in this process by developing categorical selectivity across learning in a context-dependent manner. Moreover, ACtx contains choice information and exhibits pre-stimulus biases that predict upcoming choices on ambiguous trials. These features position ACtx to route behaviorally relevant signals to downstream targets during sensory-guided behavior and illustrate how primary sensory cortices are modulated by task engagement. Together, our results clarify the differential contributions of cortical projection neuron classes and underscore the importance of ACtx in perceptual categorization and choice.

## Materials and Methods

### Mice

All procedures were approved by the University of Pittsburgh Animal Care and Use Committee and followed the guidelines established by the National Institute of Health for the care and use of laboratory animals. All mice were aged 6-10 weeks at the start of procedures and both male and female mice were included. A total of 21 wild-type C57Bl/6J mice (Jackson) were used for the L2/3 and L5 ET groups and 9 Tlx3-Cre mice (B6.FVB(Cg)-Tg(Tlx3-Cre)PL56Gsat/Mmucd, MMRRC) were used for the L5 IT group. All mice were housed under a 12 hr light/12 hr dark periodic light cycle. Mice were acclimated to head fixation for 30 min per day for 3-5 consecutive days prior to the start of behavioral or imaging experiments.

### Surgical Procedures

Mice were anesthetized with isoflurane (5%) and placed in a stereotaxic plane (Kopf model 1900). A surgical plane of anesthesia was maintained throughout the procedure via continuous delivery of isoflurane (1-2%) in oxygen. Mice lay atop a homeothermic blanket system (FHC) that maintained core body temperature at approximately 36.5^◦^C. The surgical area was shaved and cleaned with iodine and isopropyl alcohol before being numbed with lidocaine (5 mg/ml). Following all surgical procedures, antibiotic ointment was applied to wound margins, and an analgesic was administered (Carprofen, 5 mg/kg). Mice had ad libitum access to a carprofen-laced MediGel (Clear H^2^O) for 3 days post-surgery.

### Virus-mediated gene delivery

For access to L2/3 and L5 IT populations, a small incision was made on the right side of the scalp, the temporalis muscle was retracted, and two burr holes were made above ACtx lateral to the temporal ridge and approximately 1 and 2 mm rostral to the lambdoid suture. A glass pipette attached to a programmable injector (Nanoject III, Drummond) was lowered 500 *µ*m below the cortical surface and used to deliver either a non-specific GCaMP8s virus (pGP-AAV-syn-jGCaMP8s-WPRE, Addgene; titer: 2-7 x 10^12^ gc/mL) to label L2/3 neurons or a Cre-dependent GCaMP8s virus (pGP-AAV-syn-FLEX-jGCaMP8s-WPRE, Addgene; titer: 4-5 x 10^12^ gc/mL) to label L5 IT neurons. A total of 250 nL was injected into each burr hole at a rate of 10-15 nL/min (total of 500 nL). The pipette was left in place for 5-10 minutes following each injection to allow for adequate viral diffusion.

L5 ET neurons were labeled by retrograde injection into the inferior colliculus (IC). An incision was made along the midline, the skull was leveled, and a burr hole was made over right IC at coordinates 5.0 mm posterior and 0.9-1.0 mm lateral to bregma. We injected 300 nL of a retrograde GCaMP virus (pGP-AAV-syn-jGCaMP8s-WPRE, Addgene or AAV-hSyn1-GCaMP6s-P2A-nls-dTomato, Addgene; titer: 5-9 x 10^12^ gc/mL) at 2 depths (800 *µ*m and 300 *µ*m below the cortical surface) at a rate of 10-15 nL/min (total of 600 nL). The pipette was left in place for 5-10 minutes following each injection to allow for adequate viral diffusion.

### Headplate and cranial window implant

Cranial windows were created by stacking 3 round glass coverslips (3-3-4 mm, #1 thickness, Warner Instruments), which were etched with piranha solution and then bonded with transparent, UV-cured adhesive (Norland Products, Warner Instruments). The skin and periosteum above the dorsal surface of the skull was removed. To ensure strong adhesion of the dental cement, the skull surface was cleaned, dried, and treated with etchant (C&B metabond) followed by 70% ethanol. A custom headplate (iMaterialize, [130]) was affixed to the skull using dental cement (C&B metabond). A 3 mm circular craniotomy was made over right ACtx, and the cranial window was placed in the craniotomy. An airtight seal was achieved using Kwik-Sil (World Precision Instruments) around the window edges before cementing it in place. Vetbond (3M) was applied to secure the surrounding tissue to the dental cement. Implants were allowed to heal for at least 5 days before the start of experiments.

### Histological processing and anatomy

Mice were deeply anesthetized with isoflurane before transcardial perfusion with 4% paraformaldehyde in 0.01 M phosphate buffered saline. Brains were extracted and stored in 4% paraformaldehyde for 12 hr before transferring to cryoprotectant (30% sucrose) until sectioning. Sections (50 *µ*m) were cut using a cryostat (CM1950, Leica), mounted on glass slides, and coverslipped (Vectashield, Vector Laboratories). Images were obtained with an epi-fluorescence microscope (Thunder Imager Tissue, Leica) and analyzed using Fiji.

### Acoustic stimulation

Stimuli were generated with a 24-bit digital-to-analog converter (National Instruments model PXI-4461) using custom scripts written in MATLAB (MathWorks) and LabVIEW (National Instruments). Acoustic stimuli were delivered via a free-field speaker (PUI Audio) facing the left ear and calibrated using a free-field prepolarized microphone (377C01, PCB Piezotronics). Sinusoidally amplitude-modulated (sAM) broadband noise bursts (1 s duration, 5 ms cosine on/off ramps) were presented at 70 dB SPL with 100% modulation depth, spanning 2 to 32 Hz modulation frequency in 0.5 octave increments. For passive listening experiments, a 3 s inter-trial interval (ITI) separated successive sound presentations. Each stimulus was delivered in a pseudo-random order within a block, with 15-20 blocks per session.

### Mouse behavior

Prior to behavioral training, mice were water restricted for 2-3 days and provided with a daily water allotment equal to 4% of their ad libitum body weight. During training, approximately half of this water was earned through task performance, with remainder supplemented at the end of each session. Most mice stabilized at ∼80% of their ad libitum weight, and additional water was provided as needed to maintain this weight.

### Behavioral training

Mice were trained as previously described [48]. Initially, mice were trained to lick a single spout in response to either a 2 or 32 Hz sAM noise burst. The response window was 200-1500 ms after sound onset (**Fig. 1b**), and early licks were not punished. Once mice reliably licked within the response window on 10 consecutive trials, both spouts were introduced. Mice were then trained to discriminate the “easy” stimuli (2 and 2.8 Hz versus 22.6 and 32 Hz) by licking either the left or right spout. Choices were registered as the first lick within the response window. The pairing of left/right spouts with slow/fast stimuli was counterbalanced across mice. Correct responses were rewarded with a 2 *µ*L water drop, while incorrect responses resulted in a 5 s timeout. The ITI was randomly drawn from an exponential distribution ranging from 6 to 9 s. If mice developed a spout bias, defined as selecting the same spout on ≥ 6 consecutive trials, bias correction was manually implemented by presenting only the opposite-choice stimulus until mice responded correctly for 3 consecutive trials. Mice that failed to achieve *>*60% correct after 8 days of training were removed from the study.

Once mice reached a proficiency threshold of 85% correct across 200 trials, they advanced to the categorization task. In this stage, mice categorized 9 sAM noise bursts (2-32 Hz in 0.5 octave steps) as either “slow” (2 - 5.7 Hz) or “fast” (11.3 - 32 Hz) by licking the corresponding spout. The task also included a category boundary stimulus (8 Hz), which was neither slow nor fast. This boundary stimulus was presented twice within each block: once paired to the left and once to the right. Mice performed the full categorization task for 10-18 days while neural activity was recorded. At the start of each day, mice completed 20-40 easy trials to confirm that performance remained above the 85% proficiency threshold and that no response biases had emerged. If necessary, debiasing was performed until performance on easy trials returned to ≥ threshold. Debiasing was never applied to hard or boundary trials. Mice that dropped below the proficiency threshold on over half of all categorical sessions were excluded from the study.

### Licking-only task

Water-restricted mice were required to lick either the left or right spout to receive water. No auditory cues were presented, and only one spout was available per session. Each trial began after the mouse withheld licking for 3 s, which triggered delivery of a water droplet. This withholding period prevented continuous licking behavior and enabled the analysis of lick bout onsets. Once the mouse licked the spout, or if 60 s elapsed without a lick, the next trial began. Each session consisted of 30 trials. Mice required minimal shaping to produce sporadic licking in the absence of external cues.

### Two-photon imaging

Viruses were allowed to express for a minimum of 3 weeks before imaging. Widefield fluorescence imaging (Bergamo, Thorlabs) was used to measure neural responses to 4 pure tones (4, 8, 16, and 32 kHz) to generate a tonotopic map of ACtx. All two-photon fields of view (FOVs) were centered on primary auditory cortex (A1). Two-photon excitation was provided by an Insight X3 laser tuned to 940 nm (Spectra-Physics). A 16x/0.8NA water-immersion objective (Nikon) was used to acquire a 750 x 750 *µ*m, 512 x 512 pixel FOV at 30 Hz with a Bergamo II Galvo-Resonant 8 kHz scanning microscope (Thorlabs). Scanning software (Thorlabs) and stimulus generation hardware (National Instruments) were synchronized via digital pulse trains. The microscope was rotated 40-60◦ from the vertical axis to image the lateral cortex while mice remained in an upright, head-fixed position. Mice were imaged in a dark, sound attenuating chamber and monitored throughout experiments using infrared cameras (Genie Nano, Teledyne). L2/3 neurons were imaged 140-250 *µ*m below the pial surface with a power (measured at the objective) of 20-55 mW. L5 IT neurons were imaged 340-460 *µ*m below the pial surface at 30-130 mW. L5 ET neurons were imaged 400-550 *µ*m below the pial surface at 70-150 mW.

For the licking only task, we recorded multiple FOVs from L5 ET mice (*n* = 7 FOVs across 3 mice). For sAM responses in untrained mice (Naive), we collected multiple unique FOVs from each mouse (L2/3: *n* = 27 FOVs across 8 mice; L5 IT: *n* = 16 FOVs across 6 mice; L5 ET: *n* = 19 FOVs across 8 mice). With the exception of 1 mouse from each cell type, this constituted a separate experimental group from the mice used in the categorization task. For mice trained on the categorization task (L2/3: *N* = 3 mice; L5 IT: *N* = 4; L5 ET: *N* = 4), only 1 unique FOV was imaged per mouse, and the same neurons were tracked longitudinally for 10-18 days. Each imaging day began with either a passive listening block or active discrimination block, alternating across days. During passive listening blocks, the lick spouts were inaccessible to the mouse. To minimize photo bleaching, two-photon imaging during behavior was limited to a subset of trials (150-250 trials per day, lasting 30-45 minutes).

### Data Analysis

#### Behavioral quantification

Behavioral sessions were divided into early, middle, and late epochs by evenly distributing the total number of sessions for each mouse. Percent correct was calculated as the number of correct trials divided by the sum of correct and incorrect trials. Trials in which the mouse did not respond, as well as boundary trials were excluded from this calculation. Choice bias was quantified as the difference between the number of left and right choices by their sum.

Psychometric curves were fit using a cumulative Gaussian [131]:

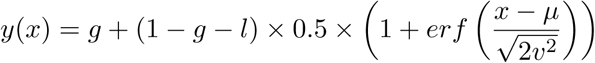

where *y*(*x*) is the probability of a rightward lick at sAM rate *x*, *erf* () is the error function, *g* is the low-frequency lapse rate, *l* is the high-frequency lapse rate (*l*), *µ* represents bias (*µ*), and *v* is a measure of discrimination sensitivity. Discrimination sensitivity (*v*) reflects the steepness of the psychometric curve (i.e., slope). However, because estimates of *v* were susceptible to noise in the presence of lapsing, psychometric slope was instead computed as the maximum derivative of the fitted psychometric function. Bias (*µ*) quantifies the *x*-axis shift of the psychometric curve relative to the category boundary (in our case, 8 Hz). Lapse rates (*g* and *l*) represent “lapses” of performance in response to easy stimuli, with the overall lapse rate quantified as *g* + *l*.

#### Processing of two-photon imaging data

Raw calcium videos were processed using Suite2P to accomplish motion correction, region of interest (ROI) detection, and spike deconvolution [132]. All ROIs were visually validated, and any low quality or non-somatic ROIs were excluded. Deconvolved activity traces were normalized by z-scoring to the mean activity in the 0.5 s period prior to stimulus onset. Across multiple imaging days, ROIs were matched longitudinally using the open-source software ROICaT (https://github.com/RichieHakim/ROICaT), and matched cells were manually validated to confirm accuracy. While some ROIs dropped out from day to day, neurons were included in the analysis if they were successfully tracked on ≥ 75% of sessions. Only 1 L5 ET mouse contained no ROIs that met this criterion.

#### Analysis of sound-evoked responses

For visualizing traces and determining sound responsiveness, activity was smoothed with a 5-frame moving average. Responses were measured within a 10-frame (333 ms) window following sound onset. A neuron was considered significantly responsive if it met two criteria: 1) ≥ 40% of the smoothed, z-scored single trial traces exceeded 0.75 standard deviations within the onset window; and 2) the trial-averaged, smoothed z-scored trace exceeded 0.5 standard deviations within the onset window. Except for analyses using matched neurons, only neurons responsive to at least 1 stimulus were included in analyses. Tuning curves were generated by averaging the activity evoked by each stimulus during the onset window. For determining significant modulation without sound (**Fig. S6**), a one-sample t-test was used to determine whether average activity within a 10-frame (333 ms) window preceding lick onset differed from a Gaussian distribution with mean of 0.

To quantify categorical selectivity, we computed a categorical selectivity index (CSI):

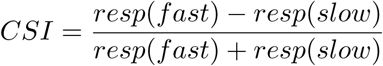

where *resp* is the average response within the onset window across all trials within a given category. For active sessions, only correct trials were included to minimize behavioral confounds. Population-level categorical selectivity was quantified as the session-average |CSI| across all neurons. To assess the bimodality of CSI distributions, we fit a 2-component Gaussian mixture model to the CSI values and measured the distance between the component means. For matched neurons that were not present in all sessions, CSI values on missing days were linearly interpolated to maintain longitudinal continuity (**Fig. S4d**).

#### Spatial clustering

To test whether categorical L5 ET neurons were spatially clustered, we measured their pairwise Euclidean distances. Neurons were classified as strongly categorical if their CSI value was *<* -0.5 or *>* 0.5. For a given category, if a FOV contained at least 5 strongly categorical neurons, we calculated the pairwise Euclidean distances among those neurons and averaged them. This value was then compared to the average pairwise distance of a random, sample-matched population within the same FOV.

#### Single-neuron decoding

We performed decoding using longitudinally matched individual neurons. A binary logistic regression classifier was used to decode slow versus fast categories on easy and hard trials, and a multinomial logistic regression classifier was used to decode stimulus identity on non-boundary trials. Decoders were trained on the average onset-window activity of each trial. For training, we extracted 10 trials of each type (either slow/fast or the 8 non-boundary stimuli) and performed 5-fold cross-validation. To ensure that all trials within a session were included in the decoding, we resampled trials with replacement 10 times. Only correct trials were used for decoding. Reported decoding accuracy for each neuron was defined as the mean accuracy of the withheld test sets across cross-validation folds and resamples. Discriminator neurons were identified as those with an average slow/fast decoding accuracy ≥ 0.8 on easy trials during early active sessions. Slow- and fast-categorical neurons were identified based on average CSI values in late active sessions (*<* -0.5 or *>* 0.5, respectively). For matrices depicting decoding accuracy across days in matched neurons (**Fig. 4, S5**), values for missing days were linearly interpolated for visualization only; interpolated values were not included in statistical analyses.

#### Generalized linear model to separate stimulus and choice

To dissociate the contributions of stimuli and choices to neural activity, we fit a generalized linear model (GLM) using the glmnet package in MATLAB (https://github.com/junyangq/glmnet-matlab). Predictor variables included stimulus identity (2-32 Hz) and lick direction (left/right). Each auditory stimulus onset contributed 10 predictors (90 total for 9 stimuli) corresponding to the first 10 imaging frames after sound onset (0–0.33 s). Licking variables (10 left and 10 right) corresponded to the 10 frames preceding lick onset (–0.33–0 s) to capture preparatory signals. All licks were included as predictors, not only those triggering reward. Time periods without predictors (i.e., no sound or licking) were excluded from the model. Predictors were standardized by dividing by their standard deviation. 2 partial models were also fit: one with only stimulus variables and one with only licking variables. In these models, the excluded variables were temporally shuffled to eliminate their relevance while preserving the number of coefficients. Each model was fit to the z-scored, deconvolved activity trace of each responsive neuron on each day. Models were trained with L1 regularization, testing 100 values of the regularization parameter *λ* and using 5-fold cross-validation to select the model with the lowest error. The coefficient of determination (*R*^2^) was calculated using predicted activity from the model:

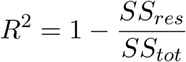

where *SS_res_* is the sum of squared residuals (the difference between observed and predicted values), and *SS_tot_*is the total sum of squares (the difference between observed values and their mean).

We used 2 complementary metrics to determine whether a neuron was driven primarily by stimulus or choice. First, we compared the *R*^2^ values of the 2 partial models 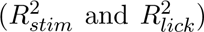. Second, we calculated the coefficient contributions of the full model:

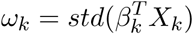

where *ω_k_* is the coefficient contribution of predictor group *k* (either stimulus or lick), *β_k_* is the vector of coefficient weights, and *X_k_* is the matrix of predictor variables. This normalization allowed us to compare weights across unequal numbers of coefficients (90 stimulus predictors vs 20 lick predictors). A given neuron was labeled as a “stimulus” neuron if 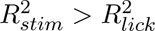 and *ω_stim_ > ω_lick_* or labeled as a “choice” neuron if 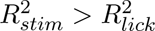 and *ω_stim_ < ω_lick_*. The small fraction of neurons with conflicting outcomes (e.g. 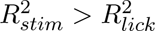 but *ω_stim_ < ω_lick_*) between the 2 test metrics were excluded from either group.

#### Clustering

For each cell type, we performed hierarchical clustering on the responses of stimulus and choice neurons. Within each cell type, neurons were concatenated across sessions for each subgroup. For each neuron, we calculated the mean onset responses on left and right trials, separated by correctness. This produced 4 response values per neuron (correct left, correct right, incorrect left, incorrect right). Responses were rescaled to a 0-1 range and clustered using Euclidean distance with Ward’s linkage methods (MATLAB linkage and cluster functions) to generate dendrograms. The number of clusters was manually curated based on the dendrogram, distance matrix, and underlying neural responses, selecting the fewest clusters necessary to capture the heterogeneity of response types.

#### Population decoding

To quantify how neural populations encoded task variables across time and learning, we applied an L2-regularized logistic regression decoder (scikit-learn LogisticRegression, default parameters) on a per-session, per-frame basis. Decoding was performed separately on data aligned to stimulus onset or lick onset. Sessions with fewer than 10 responsive neurons (including stimulus and choice subgroups) were excluded. For each classifier, we randomly sampled 100 neurons with replacement and balanced the number of slow and fast (or left and right) trials via downsampling. Both correct and incorrect trials were included for decoding. Classifiers were trained using 5-fold cross validation. Neurons and trials were resampled 20 times, and the reported decoding accuracy for each session and frame was the mean test-set accuracy across all cross validation folds and resamplings. Chance performance was estimated by independently shuffling trial labels within each decoder resampling.

#### Dimensionality reduction

To visualize time-varying population activity on boundary trials, we used principal component analysis (PCA) to reduce the dimensionality of our dataset. For our example session (**Fig. 7a**), activity from individual trials was projected onto the first 3 principal components (PCs), revealing separation between left- and right-choice trials. We then utilized a more rigorous PCA procedure that was capable of quantifying this effect across sessions. We began by randomly sampling 100 neurons with replacement and balancing left and right boundary-choice trials via downsampling.

Sessions were included only if they contained at least 5 boundary trials of each choice and at least 10 responsive neurons (including stimulus and choice subgroups). For each qualifying session, we computed the mean responses for left and right choices on boundary trials (aligned either to stimulus onset or to lick onset) and performed PCA on these data. The first 10 PCs were extracted, and the Euclidean distance between the 2 mean trajectories was computed to obtain a time-resolved measure of trajectory separation. This procedure was repeated 20 times with resampling of neurons and trials (with replacement), and the reported distance for each session was the average separation across all resamplings.

To test whether pre-stimulus activity could predict upcoming choices on the perceptually ambiguous boundary stimulus, we applied PCA analogous to the procedure described above. For each session, PCA was performed exclusively on correct non-boundary trials. PC-projected activity from these trials at lick onset was averaged separately for left and right choices to compute centroids for “choose left” and “choose right”. On boundary trials, we extracted the population activity immediately preceding stimulus onset and calculated its Euclidean distance to each choice centroid. The upcoming choice was predicted as left or right based on the closer centroid in PC space, and this prediction was compared to the true outcome to yield a prediction accuracy per session. To generate a null distribution for statistical comparison, we shuffled the true left/right labels of boundary trials and recomputed prediction accuracy on the shuffled data.

### Statistical analysis

All statistical analyses were performed in MATLAB (Mathworks) and Prism (Graphpad). Data are reported as mean +*/*− SEM unless otherwise stated. In cases where the same data sample was used for multiple comparisons, all p-values remained significant after correction (\**p <* 0.05, \*\**p <* 0.01, \*\*\**p <* 0.001, \*\*\*\**p <* 0.0001).

## Acknowledgments

We thank current and former members of the Williamson Lab for helpful feedback and assistance with animal care. Special thanks to Joshua John and Jocelyn Martin for assistance with behavioral training and Megan Arnold for assistance with surgeries. This work was supported by NIH/NIDCD grants R21DC018327784 and R01DC020459, a Hearing Health Foundation Emerging Research Grant, and the Klingenstein-Simons Fellowship in Neuroscience to RSW. Further support was provided by NIH/NIDCD grant F31DC021363 (NAS), NRSA grant T32NS007433 (NAS), and NIH grant T32NS126122 (MM).

## Author Contributions

NAS and RSW conceptualized all experiments. NAS collected all data. NAS and MM analyzed all data. NAS and RSW prepared figures and wrote the manuscript. NAS, MM, and RSW reviewed and edited the manuscript.

---

**Figure S1:**
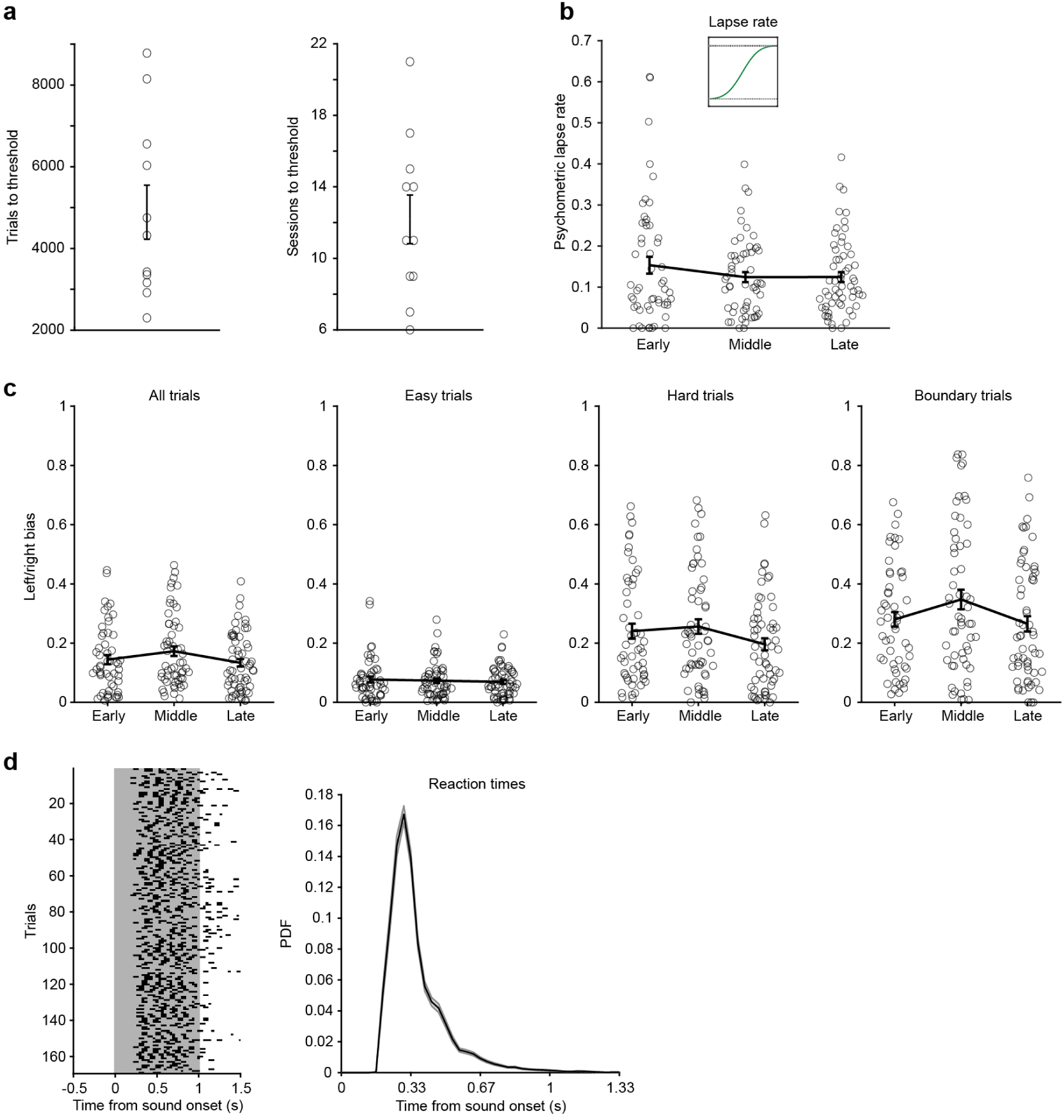
Lapse rate and choice bias do not change with learning. (a) The number of trials and sessions it took for each mouse to reach the proficient threshold necessary to begin imaging. (b) Lapse rate of psychometric curves fit to each behavioral session broken up by learning phase (one-way ANOVA, main effect for learning, *p* = 0.30; early: *n* = 53 sessions, middle: *n* = 57 sessions, late: *n* = 60 sessions; *N* = 11 mice). (c) Choice bias for each session across learning within all trials (one-way ANOVA, main effect for learning, *p* = 0.15), easy trials (one-way ANOVA, main effect for learning, *p* = 0.78), hard trials (one-way ANOVA, main effect for learning, *p* = 0.16), and boundary trials (one-way ANOVA, main effect for learning, *p* = 0.09; early: *n* = 53 sessions, middle: *n* = 57 sessions, late: *n* = 60 sessions; *N* = 11 mice). (d) Left: a licking raster across trials for an example session. Right: average probability density function of reaction times.

**Figure S2:**
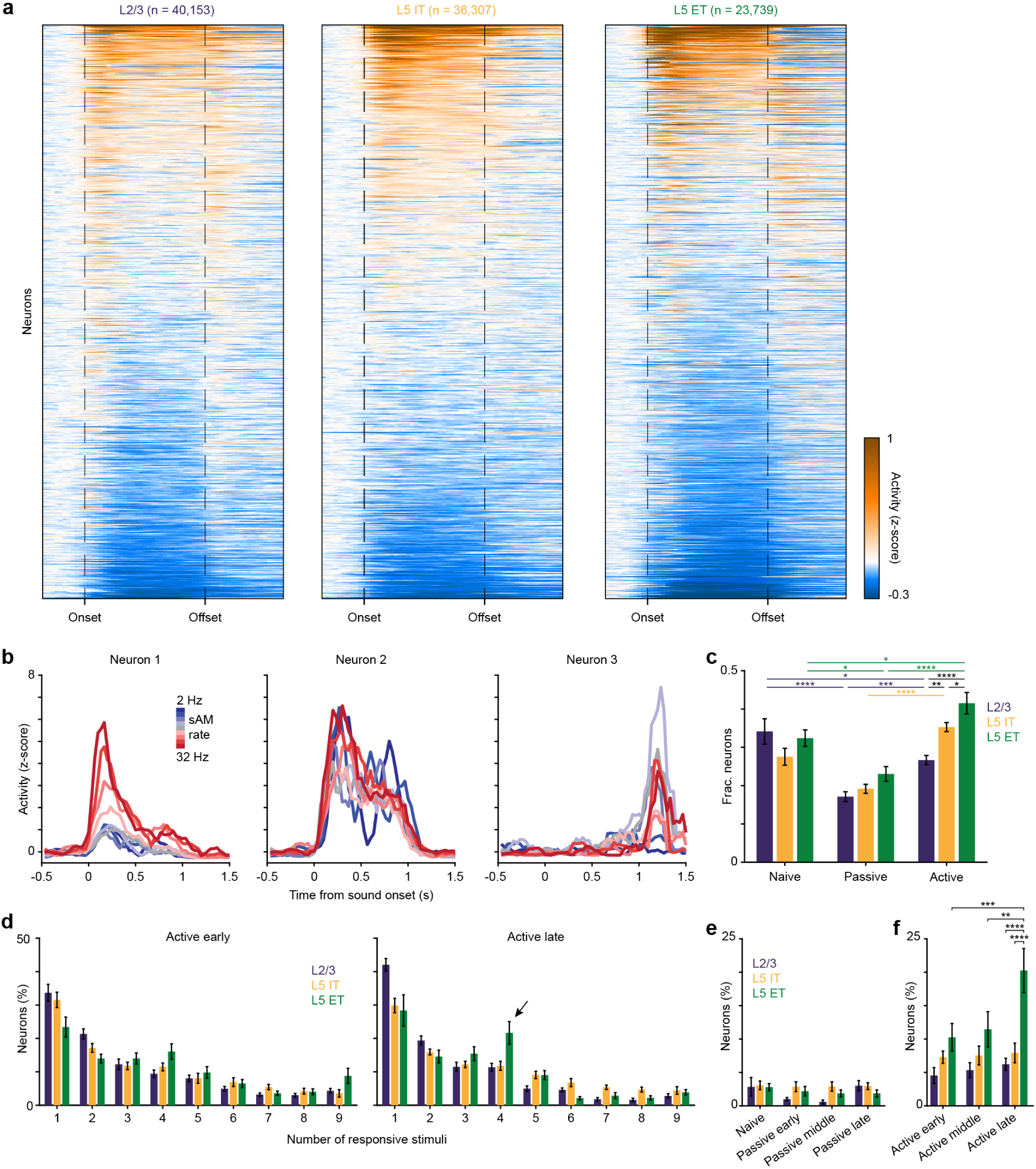
Heterogeneity of sound responses across cell types. (a) Trial-averaged responses of all recorded neurons for each cell type sorted by average stimulus-evoked activity. (b) Average stimulus responses for an example onset, sustained, and offset neuron. (c) Proportion of onset responsive neurons across cell types and contexts (L2/3 naive: *n* = 27 sessions, L5 IT naive: *n* = 16, L5 ET naive: *n* = 19, L2/3 passive: *n* = 49, L5 IT passive: *n* = 63, L5 ET passive: *n* = 53, L2/3 active: *n* = 49, L5 IT active: *n* = 63, L5 ET active: *n* = 58; Two-way ANOVA with post-hoc Tukey test, main effect for cell type, *p* = 0.0010, main effect for context, *p <* 0.0001, interaction, *p* = 0.0018, asterisks represent pairwise comparisons). (d) Distribution of neurons based on how many sAM stimuli elicit a significant response in early and late active behavior. The arrow emphasizes an increase in L5 ET neurons responding to four stimuli late in learning. (e) Percentage of neurons responding selectively to the four slow or four fast categorical stimuli in naive and passive listening. (f) Same as (e) for active behavior (L2/3 active early: *n* = 15, L5 IT active early: *n* = 20, L5 ET active early: *n* = 18, L2/3 active middle: *n* = 17, L5 IT active middle: *n* = 21, L5 ET active middle: *n* = 19, L2/3 active late: *n* = 17, L5 IT active late: *n* = 22, L5 ET active late: *n* = 21; Two-way ANOVA with post-hoc Tukey test, main effect for cell type, *p <* 0.0001, main effect for learning, *p* = 0.02, asterisks represent pairwise comparisons).

**Figure S3:**
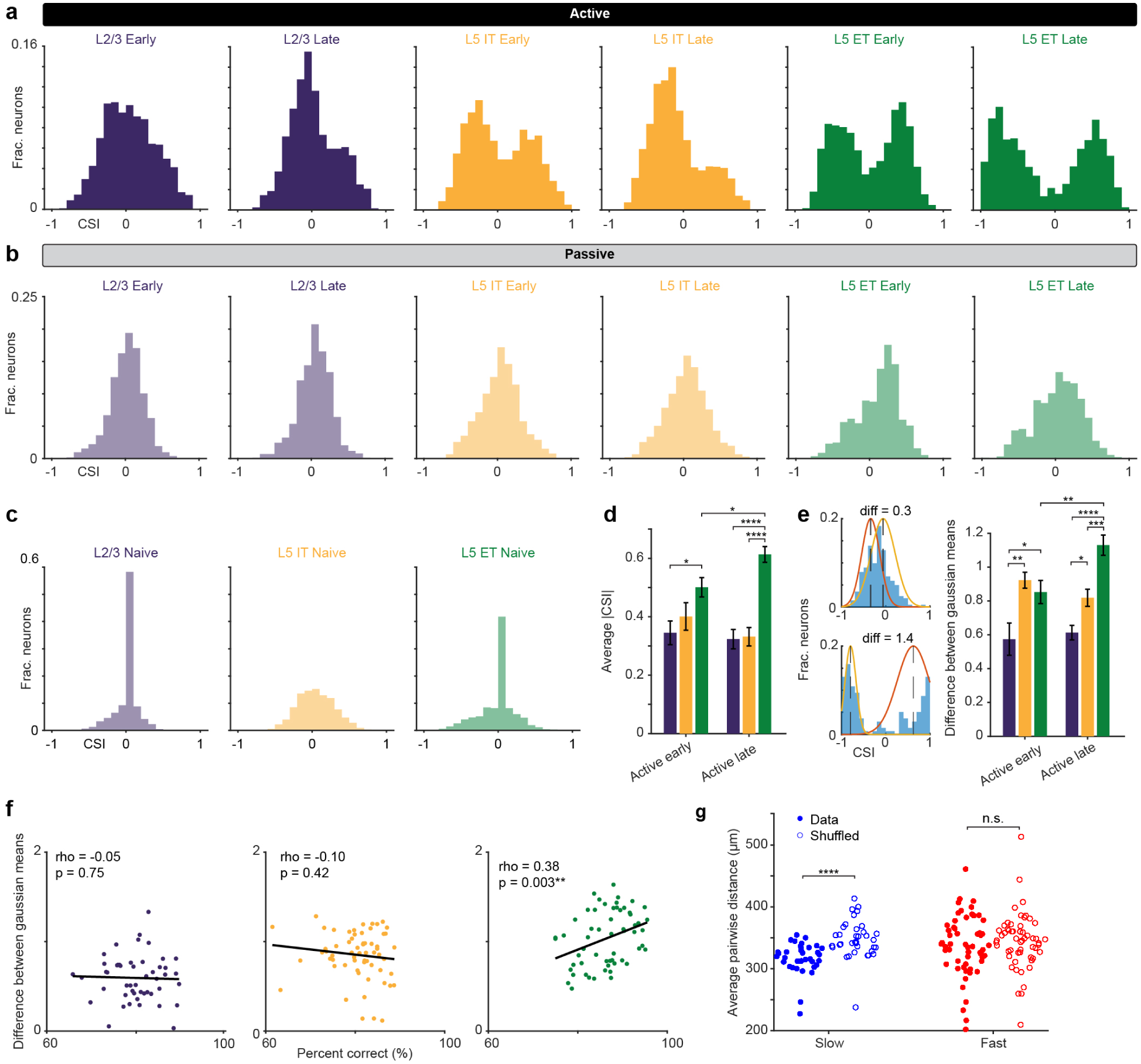
Categorical selectivity index shows cell-type-specific effects. (a) Histograms of CSI values on all trials for neurons responsive during active behavior. Plots are separated by early and late sessions across cell types. (b) Histograms of CSI values on all trials for neurons responsive during passive listening. (c) Histograms of CSI values on all trials for neurons responsive during naive sessions. (d). Average absolute value of CSIs calculated with correct trials only for active sessions broken down by learning and cell type (L2/3 early: *n* = 15 sessions, L5 IT early: *n* = 20, L5 ET early: *n* = 18, L2/3 late: *n* = 17, L5 IT late: *n* = 22, L5 ET late: *n* = 21; Two-way ANOVA with post-hoc Tukey test, main effect for cell type, *p <* 0.0001, interaction, *p* = 0.02, asterisks represent pairwise comparisons). (e) Left: Examples of fitting a two-component gaussian model to a CSI histogram and the difference of the means. Right: Difference of gaussian means of CSI histograms calculated with correct trials only for active sessions broken down by learning and cell type (L2/3 early: *n* = 15 sessions, L5 IT early: *n* = 20, L5 ET early: *n* = 18, L2/3 late: *n* = 17, L5 IT late: *n* = 22, L5 ET late: *n* = 21; Two-way ANOVA with post-hoc Tukey test, main effect for cell type, *p <* 0.0001, interaction, *p* = 0.003, asterisks represent pairwise comparisons). (f) Correlation between the difference of gaussian means and the percent of correct trials on each session. L5 ET shows a significant correlation (L2/3: *n* = 49 sessions, L5 IT: *n* = 63, L5 ET: *n* = 58; Spearman’s Rho). (g) Average pairwise distances of slow and fast L5 ET categorical neurons within an imaging plane compared to a shuffled distribution. Slow categorical neurons show significant spatial clustering compared to chance (slow: *n* = 36 sessions, paired t-test, *p <* 0.0001; fast: *n* = 56 sessions, paired t-test, *p* = 0.37)

**Figure S4:**
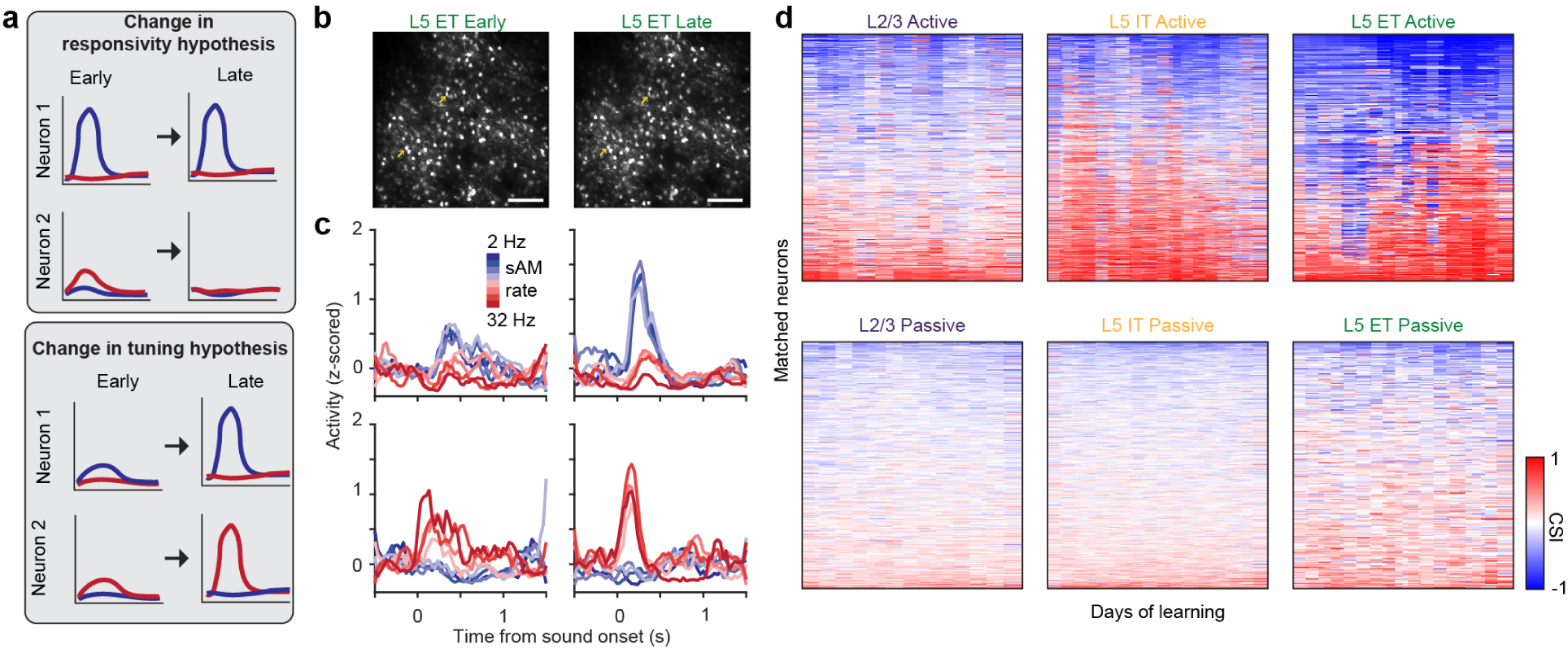
Individual L5 ET neurons dynamically shift responses to encode categories. (a) Two hypotheses for how categorical selectivity could arise at the population level. (b) Example L5 ET field of view matched between early and late in learning. Arrows show examples in (c) matched across FOVs. Scale bar represents 150*µm*. (c) Example stimulus responses from neurons matched from early to late in learning illustrating the development of categorical selectivity. (d) Matrix of CSI values for all neurons matched longitudinally across all days of training. Each row is a unique neuron and colors denote the CSI value. Neurons are sorted by average CSI value across all days.

**Figure S5:**
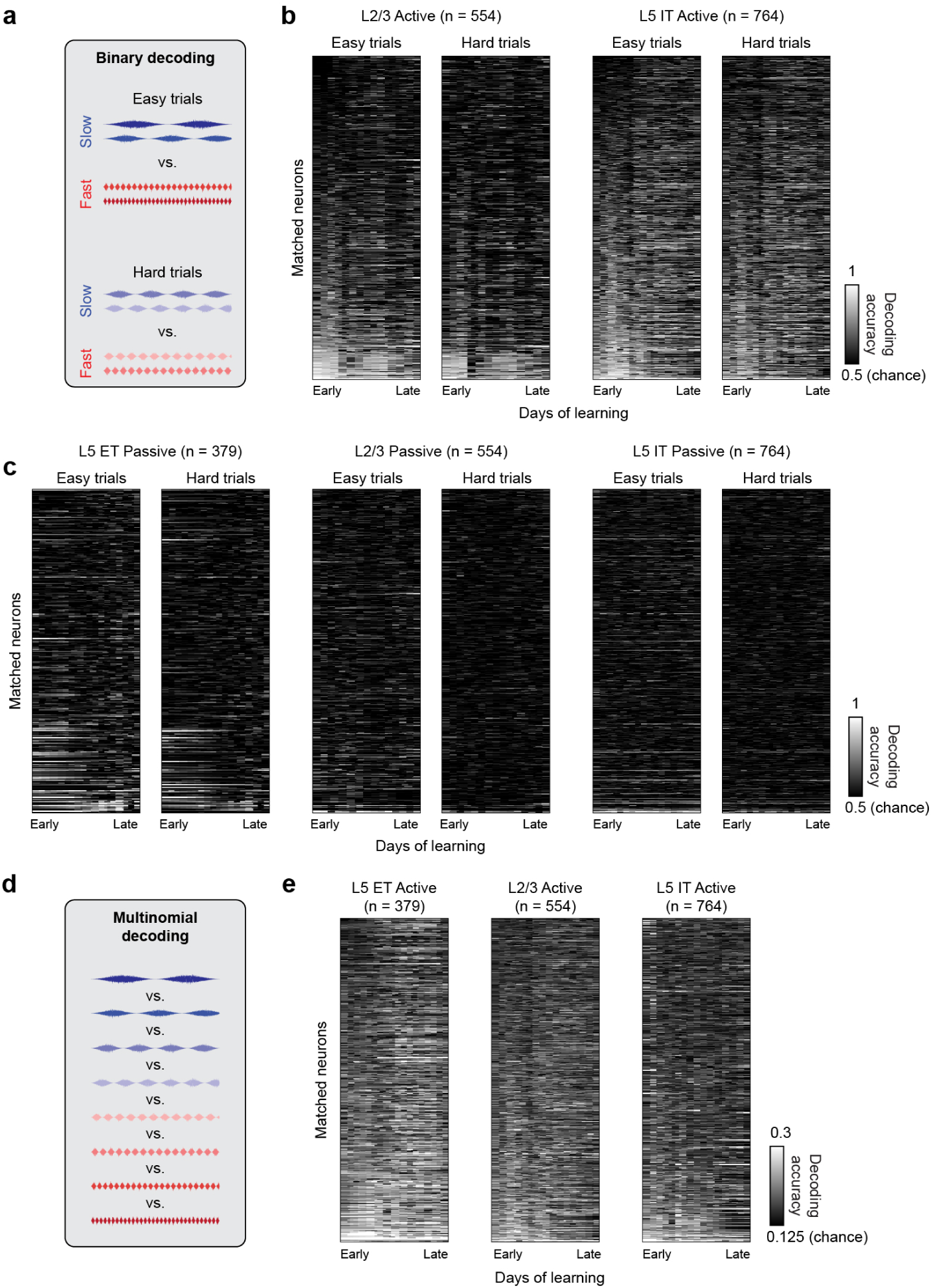
Single neuron decoding reveals discriminator neurons. (a) Schematic of binary decoding for easy and hard trials. (b) Binary decoding accuracies for matched L2/3 and L5 IT neurons across learning in active sessions for both easy and hard stimuli. (c) Binary decoding accuracies for matched L5 ET, L2/3, and L5 IT neurons across learning in passive sessions for both easy and hard stimuli. (d) Schematic for multinomial decoding of stimulus identity. (e) Stimulus decoding accuracies for matched L5 ET, L2/3, and L5 IT neurons across learning in active sessions.

**Figure S6:**
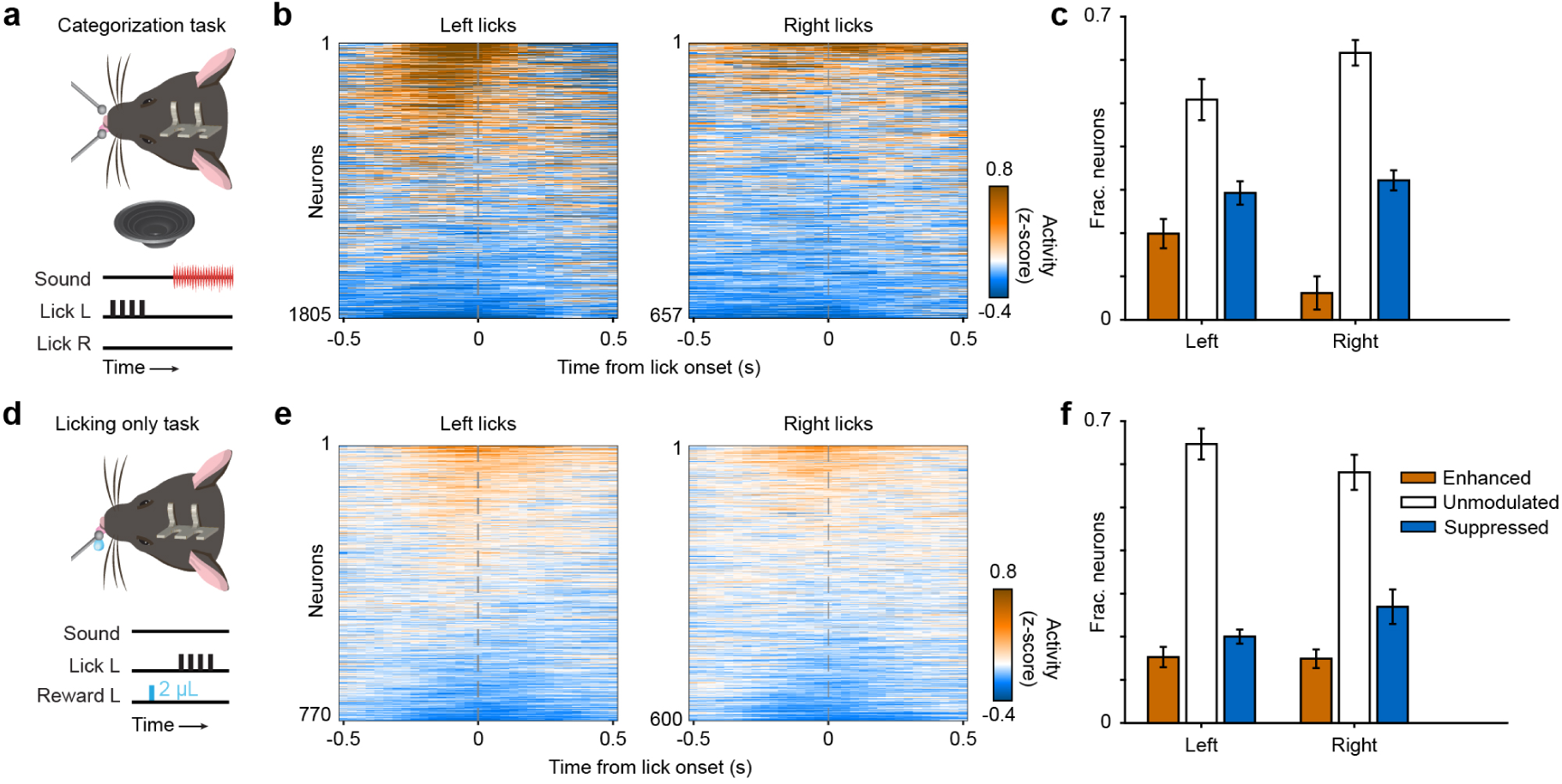
L5 ET neurons are modulated by movement. (a) Schematic of the categorization task focusing on licking activity prior to the onset of sound. (b) L5 ET neuron activity averaged across trials where mice licked before the presentation of sound. (c) Average fraction of L5 ET neurons significantly modulated by licking activity alone in the categorization task. (d) Schematic of a licking only task where mice periodically lick a spout to receive water without any auditory cues. (e) L5 ET neuron activity averaged across trials where mice licked to collect water. (f) Average fraction of L5 ET neurons significantly modulated by licking activity in a licking only task.

**Figure S7:**
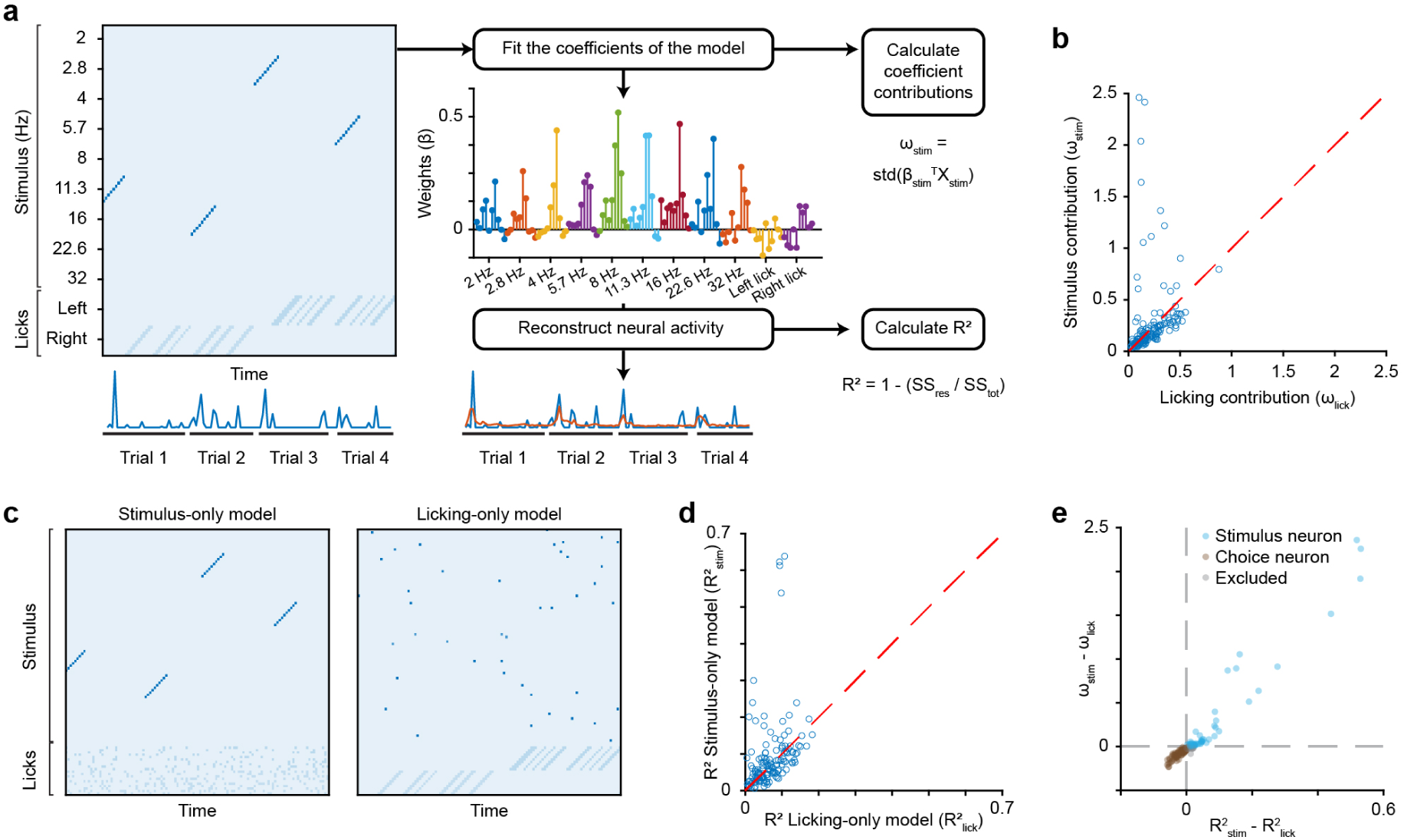
A generalized linear model can separate stimulus and choice subpopulations. (a) Schematic of how to fit a generalized linear model and extract relevant information. First, a matrix of behavioral variables is assembled and aligned with the activity of a single neuron. Next, the coefficients of the model are fit to the neural activity. The contributions of coefficient groups (stimulus and licking) can then be calculated. Finally, the coefficients are used to reconstruct the neural activity and the coefficient of determination is calculated by comparing the predicted activity to the true activity. (b) Comparison between coefficient contributions (*ω*) for all neurons in an example session. (c) Example design matrices (*X*) for two partial models containing only stimulus or licking information. (d) Comparison between *R*^2^ values of the two partial models for all neurons in an example session. (e) Comparison between *ω* and partial model *R*^2^ within an example session to illustrate how neurons are classified as either a stimulus or choice neuron.

**Figure S8:**
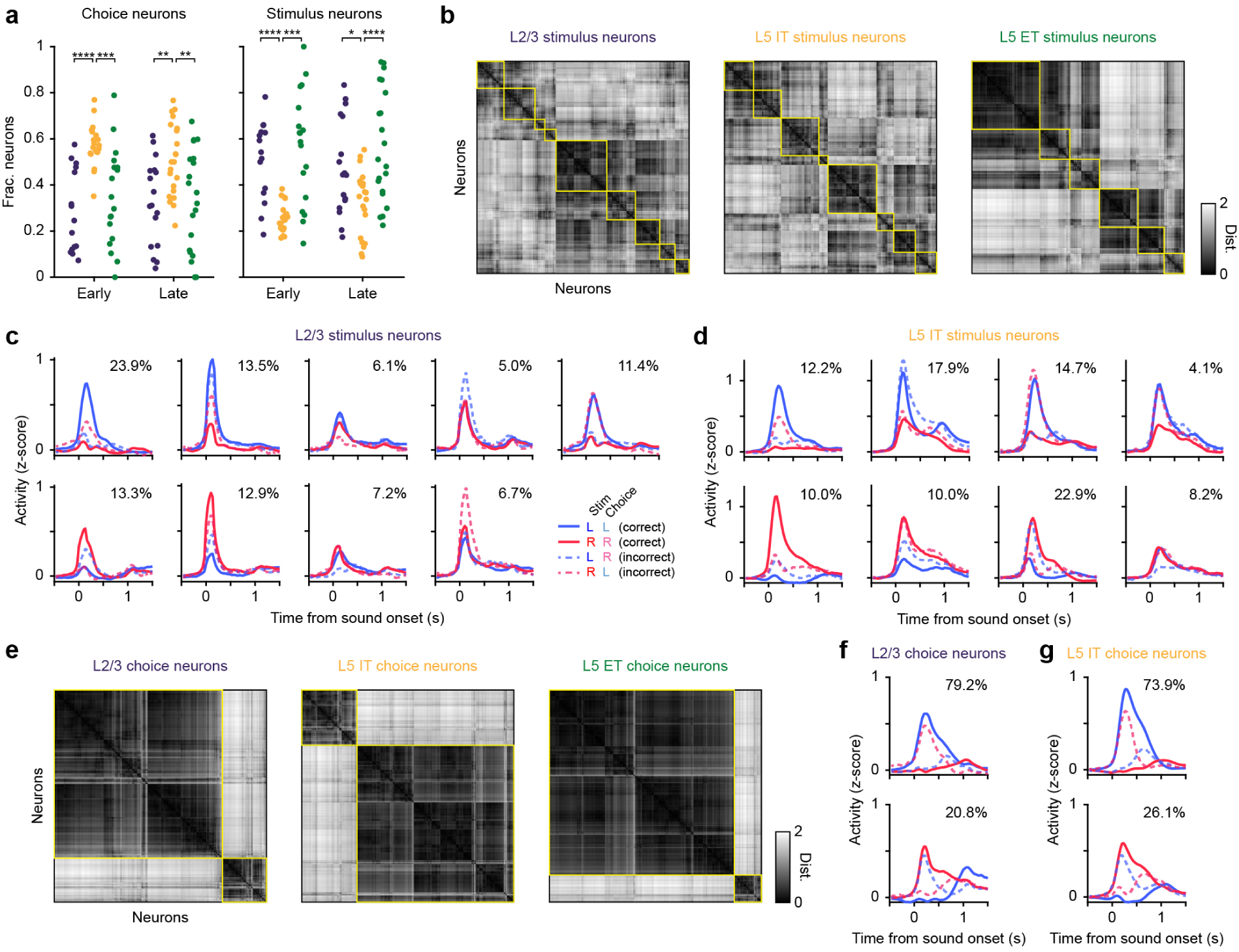
Stimulus and choice neurons across cell types. (a) Fraction of stimulus and choice neurons across cell types in early and late behavior (L2/3 early: *n* = 15 sessions, L5 IT early: *n* = 20, L5 ET early: *n* = 18, L2/3 late: *n* = 17, L5 IT late: *n* = 22, L5 ET late: *n* = 21; Stimulus neurons: Two-way ANOVA with post-hoc Tukey test, main effect for cell type, *p <* 0.0001, Tukey test for multiple comparisons; Choice neurons: Two-way ANOVA with post-hoc Tukey test, main effect for cell type, *p <* 0.0001, asterisks represent pairwise comparisons). (b) Distance matrices for hierarchical clustering of stimulus neurons across cell types. Yellow outlines denote clusters. (c) Average cluster responses for L2/3 stimulus neurons. Percentage represents the respective size of each cluster within the population. (d) Same as (c) for L5 IT stimulus neurons. (e) Same as (b) for choice neurons. (f) Same as (c) for L2/3 choice neurons. (g) Same as (c) for L5 IT choice neurons.

**Figure S9:**
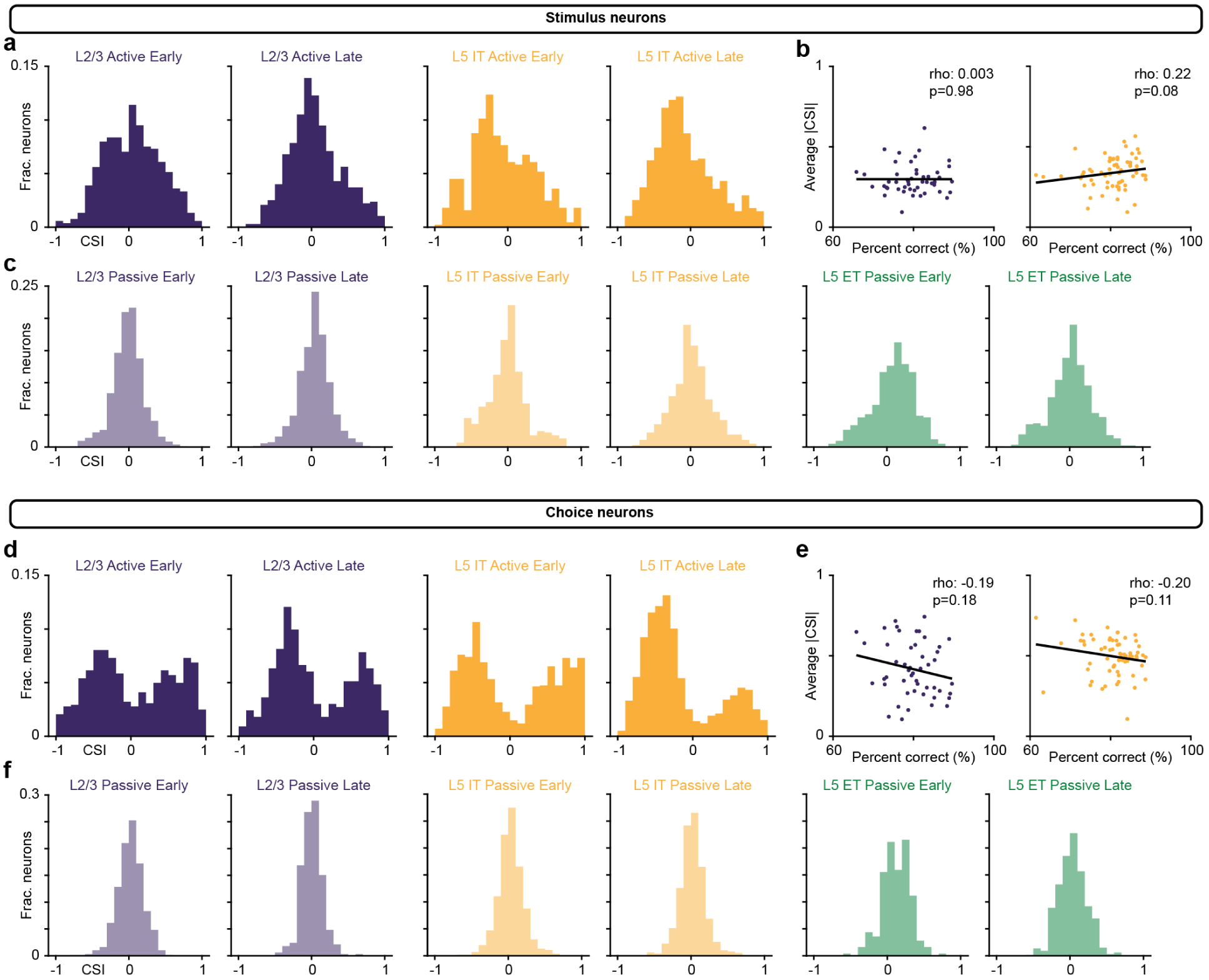
Categorical selectivity does not change across learning in L2/3 or L5 IT stimulus or choice neurons. (a) CSI histograms for stimulus neurons during behavior. (b) Correlation between average absolute value CSIs and percent correct in each session. Line represents linear regression fit (L2/3: *n* = 49 sessions, L5 IT: *n* = 63; Spearman’s Rho). (c) CSI histograms for stimulus neurons during passive listening. (d) Same as (a) for choice neurons. (e) Same as (b) for choice neurons (L2/3: *n* = 49 sessions, L5 IT: *n* = 63; Spearman’s Rho). (f) Same as (c) for choice neurons.

**Figure S10:**
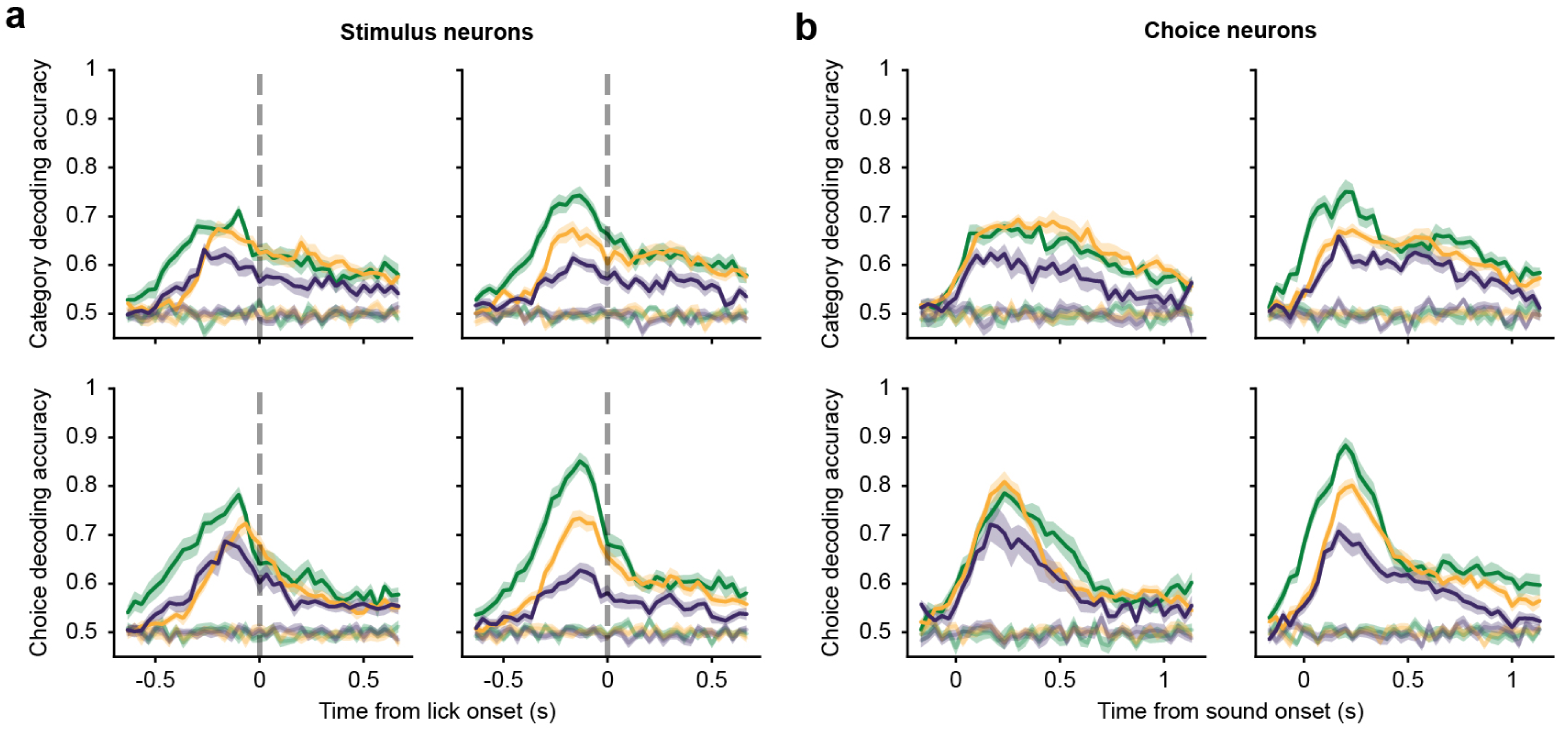
Population decoding in functional subpopulations. (a) Average category (top) and choice (bottom) decoding accuracies for stimulus neurons across the duration of a trial aligned to lick onset. Transparent lines represent shuffled data. (b) Same as (a) for choice neurons aligned to stimulus onset.

**Figure S11:**
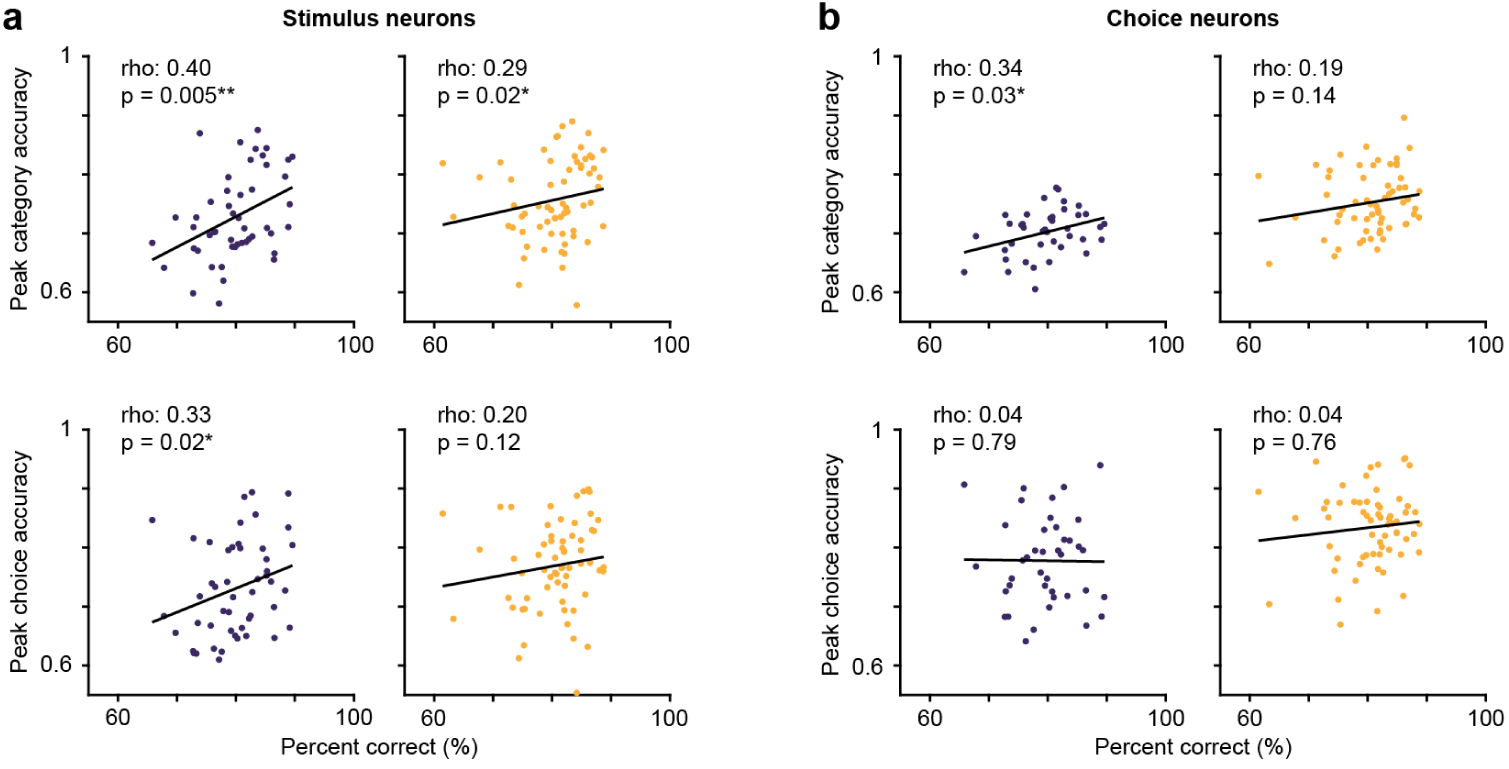
Correlations between behavioral performance and decoding accuracy in L2/3 and L5 IT neurons. (a) Average category (top) and choice (bottom) decoding accuracies for stimulus neurons correlated with performance in each session. Lines represent linear regression fit (L2/3: *n* = 49 sessions; L5 IT: *n* = 63; Spearman’s Rho). (b) Same as (a) for choice neurons (L2/3: *n* = 40 sessions; L5 IT: *n* = 62; Spearman’s Rho).

**Figure S12:**
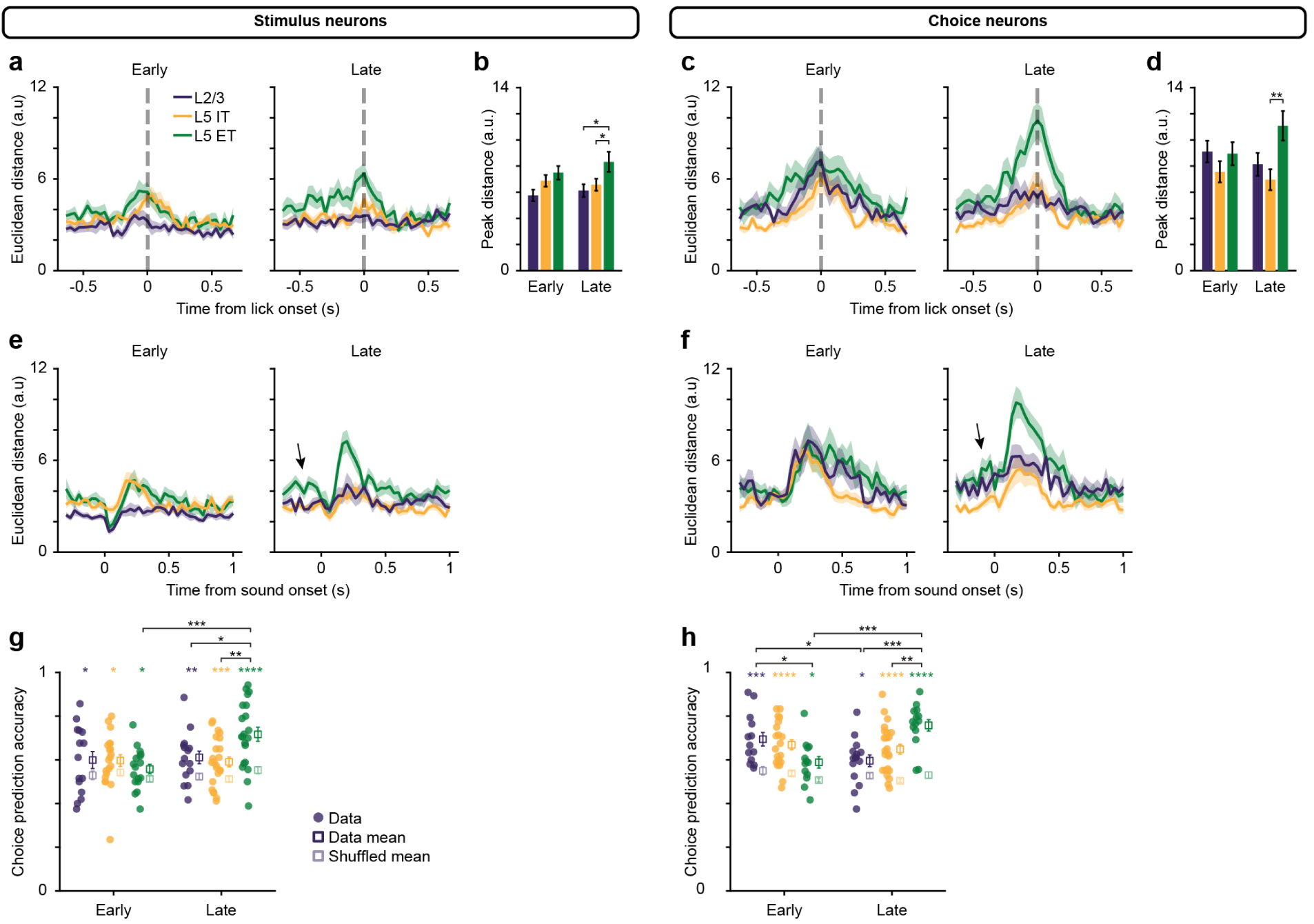
Encoding of ambiguous stimuli in stimulus and choice neurons. (a) Distance between average lick-aligned left and right trajectories for stimulus neurons. (b) Peak distance between lick-aligned trajectories for stimulus neurons across early and late sessions (L2/3 early: *n* = 15 sessions, L5 IT early: *n* = 20, L5 ET early: *n* = 17, L2/3 late: *n* = 15, L5 IT late: *n* = 23, L5 ET late: *n* = 21; Two-way ANOVA with post-hoc Tukey test, main effect for cell type, *p* = 0.004, asterisks represent pairwise comparisons). (c) Same as (a) for choice neurons. (d) Same as (b) for choice neurons (L2/3 early: *n* = 13 sessions, L5 IT early: *n* = 20, L5 ET early: *n* = 13, L2/3 late: *n* = 15, L5 IT late: *n* = 23, L5 ET late: *n* = 14; Two-way ANOVA with post-hoc Tukey test, main effect for cell type, *p* = 0.009). (e) Distance between average stimulus-aligned left and right boundary trial trajectories for stimulus neurons. Black arrow emphasizes pre-stimulus choice information in L5 ET neurons. (f) Same as (e) for choice neurons. (g) Accuracy of predicting a boundary choice based on the pre-stimulus population activity in stimulus neurons (L2/3 early: *n* = 15 sessions, paired t-test, *p* = 0.03; L5 IT early: *n* = 20, paired t-test, *p* = 0.02; L5 ET early: *n* = 17, paired t-test, *p* = 0.04; L2/3 late: *n* = 15, paired t-test, *p* = 0.001; L5 IT late: *n* = 23, paired t-test, *p* = 0.0004; L5 ET late: *n* = 21, paired t-test, *p <* 0.0001, colored asterisks represent comparisons against shuffled data; Two-way ANOVA with post-hoc Tukey test, main effect for learning *p* = 0.03, interaction *p* = 0.01, black asterisks represent pairwise comparisons between groups). (h) Same as (g) for choice neurons (L2/3 early: *n* = 13 sessions, paired t-test, *p* = 0.0002; L5 IT early: *n* = 20, paired t-test, *p <* 0.0001; L5 ET early: *n* = 13, paired t-test, *p* = 0.02; L2/3 late: *n* = 15, paired t-test, *p* = 0.01; L5 IT late: *n* = 23, paired t-test, *p <* 0.0001; L5 ET late: *n* = 14; paired t-test, *p <* 0.0001, colored asterisks represent comparisons against shuffled data; Two-way ANOVA with post-hoc Tukey test, interaction *p >* 0.0001, black asterisks represent pairwise comparisons between groups).

